# Age- and Virus-Specific Signatures of *In Vitro* Reconstituted Human Airway Epithelia in the Presence and Absence of Respiratory Viral Infections

**DOI:** 10.1101/2025.02.16.638411

**Authors:** Mathilde Bellon, Catia Alvarez, Gustavo Ruiz Buendia, Pascale Sattonnet-Roche, Damien Dbeissi, Yoann Sarmiento, Laurent Kaiser, Arnaud Didierlaurent, Isabella Eckerle, Manel Essaidi-Laziosi

**Affiliations:** Department of Medicine, Faculty of Medicine, University of Geneva; Geneva Centre for Emerging Viral Diseases, Geneva University Hospitals and University of Geneva; Translational Data Science, SIB Swiss Institute of Bioinformatics; Division of Infectious Diseases, Geneva University Hospitals; Centre for Vaccinology, Department of Pathology and Immunology, University of Geneva

**Keywords:** RSV, SARS-CoV-2, RV, influenza virus, human airway epithelia, virus-host interaction, age-specific, innate immune response, muco-ciliary clearance

## Abstract

While Influenza Virus and Respiratory Syncytial Virus (RSV) are considered as a significant health burden in children, Severe Acute Respiratory Syndrome Coronavirus-2 (SARS-CoV-2) causes milder diseases in this age group compared to adults. To investigate the involvement of the upper respiratory tract human airway epithelium (HAE) in this pattern, we established an in-house model of reconstituted HAE cultured in air-liquid interface from nasal swabs of children and adults and characterised it before and after *ex vivo* respiratory viral infections using focused and unbiased approaches. Fully differentiated paediatric HAE exhibited an increasing induction level of genes related to mucociliary clearance, while higher expression of innate immune pathways was found in the ones from adults. While similar viral replication kinetics in both age groups were shown for SARS-CoV-2, Influenza A Virus (IAV), RSV and Rhinovirus (RV) infection, transcriptomic analysis showed stronger and earlier induction of IFN-related pathways in SARS-CoV-2-infected HAE from children compared to IAV, RSV and RV. IAV and RSV had the weakest innate immune response increase in HAE from children versus adults. RV infection showed an intermediate pattern, resembling SARS-CoV-2 more than RSV or IAV. Our work demonstrates a distinct sensing of SARS-CoV-2 compared to other respiratory viruses *ex vivo*, which may contribute to the milder course of disease in SARS-CoV-2 infected children and argues for a role of early virus-HAE interaction in shaping viral pathogenesis. Furthermore, we show that innate immune responses towards respiratory viruses are virus-specific and differ between age groups. Hence, findings on SARS-CoV-2 cannot be extrapolated to other respiratory viruses.

## Introduction

Viral respiratory tract infections are among the most common diseases in humans^1^, caused by a wide range of viruses from different families. Respiratory viruses can infect the upper and/or lower respiratory tract and display a broad clinical spectrum varying from asymptomatic to mild or moderate disease, to pneumonia and life-threatening disease, especially in vulnerable groups. Respiratory viral infections constitute a global public health concern and have a significant socio-economic impact.

Children have the highest prevalence of viral respiratory tract infections^2^, with an average of 9 episodes per year and per child^3,4^. The most prevalent virus and the typical causative agent of the common cold in children are human Rhinoviruses (RV), while the most relevant in terms of morbidity are Influenza Virus and Respiratory Syncytial Virus (RSV). Since 2019, a novel virus, Severe Acute Respiratory Syndrome Coronavirus-2 (SARS-CoV-2), the etiological agent of Coronavirus Disease 2019 (COVID-19), has added to the continuously circulating respiratory viruses, with repeated infection waves.

The clinical manifestations of these viruses differ according to age. For instance, while RSV constitutes the leading cause of acute lower respiratory infection in children younger than 5 years and the primary cause of bronchiolitis in infants^5^, it mainly causes subclinical/mild infections in healthy adults. Influenza Virus presented the first cause of hospitalization for acute respiratory tract illness in school-aged children before the emergence of SARS-CoV-2^6^. Children are also part of the Influenza at-risk population with an increasing risk of disease severity during the first years of life^7^. On the contrary, despite SARS-CoV-2 global impact and significant morbidity and mortality in adults, on average, has presented less severe in children, with mostly asymptomatic or mild acute presentation^8^.

The differences in disease presentation between viruses and age groups remain only partly understood. Several studies have previously demonstrated an age-dependent induction of innate and adaptive immunity. The study of cytokines induction and immune cell activation had previously pointed out the implication of an unbalanced type 1 and type 2 immune responses in Influenza- and RSV-mediated acute lower respiratory tract infections and bronchiolitis in children, leading to an inadequate anti-viral response^9–12^. In contrast, comparative transcriptomic analyses of epithelial and immune cells from clinical nasal swabs have more recently shown better efficacy of the early antiviral state induction against SARS-CoV-2, which is sensitive to interferon (IFN), in children and more transient immune cells’ activation, including neutrophils and dendritic cells. This likely leads to a more vigorous host response and faster return to homeostasis, presumably responsible for a lighter and shorter duration of COVID-19 symptoms^13–15^.

These clinical data hint towards an age-dependent host response during acute viral infections and suggest distinct viral sensing by the airway epithelia at the early course of viral infection in children. An in-depth examination of the involvement of the airway epithelium intrinsic response, as the first infection hub and physicochemical barrier against pathogens (reviewed in^16^), in modulating clinical variability is still needed to better understand respiratory viral pathogenesis in children compared to adults. The goal of our study was, therefore, to investigate innate immune responses in an *ex vivo* model of the nasal human airway epithelium (HAE) in children versus adults at baseline and during acute infection with SARS-CoV-2, Influenza A Virus (IAV), RV and RSV. For this purpose, we first successfully established an in-house *in vitro* differentiated HAE model reconstituted from nasal swabs collected from children and adults. After the morphological and functional characterization of the polarized tissue, we assessed viral replication by quantification of viral RNA and the innate immune response by RNA sequencing before and during virus infection.

## Material and methods

### Participant inclusion and human material collection

In this study, approved by the Geneva Ethics Committee (CCER project number 2022-01601), we included children between 3 and 10 years old and adults (> 18 years old), summarized in Supplementary TableS1, without known respiratory comorbidities, immunological or coagulation disorders, acute/chronic infections, immunomodulator and/or anticoagulant treatment. In case of anamnestically reported acute respiratory tract symptoms, the sample collection was postponed until the symptoms resolved. Before sampling, written informed consent was obtained from adult participants and children’s legal representatives, while verbal assent to participate was obtained from children after an age-adapted explanation of the study. After a nasal wash with a saline solution (Physiodose, Laboratoires Gilbert), nasal airway epithelial cells were collected with a nasal swab by gently inserting a cytological brush (Combiplus cell collector, Trimastek Medial) in the nostrils of the participant and rotating it over the inferior turbinate during approximatively 5 seconds. Both nostrils were sampled. The swabs were directly placed in a tube containing culture medium for isolation (PneumaCult Ex+ Medium, (StemCell) with 2% of Penicilline/Streptomycine (Gibco by Life Technologies) and 0.5% of Amphotericin B (Gibco by Life Technologies)) and then immediately transferred to the laboratory for further processing. In total, samples were collected from 17 children and 20 adults, including 9 families (children and parents) and 3 adults unrelated to any children.

### Primary nasal epithelium isolation, expansion, and differentiation

#### 1. Cell isolation

Twelve-well plates were pre-coated with a coating solution of RPMI (Gibco) containing 30μg/mL collagen I rat tail (Gibco), 10μg/mL of human fibronectin (Sigma) and 10μg/mL of bovine serum albumin (BSA, Sigma), incubated at least 3 hours (h) at 37°C and then washed 3 times with phosphate buffered saline (PBS, 1x without calcium, without magnesium pH 7.4, Gibco). The collected cells were seeded in the pre-coated wells after the brushes were scratched between them or with a pipet tip to detach the cells into the isolation medium. In case of slight blood contamination during the collection due to nose bleeding, blood was gently removed from the brush. If not possible, the blood-containing sample was excluded. Cells collected from the nasal swabs were seeded in 2 wells per donor. They were then incubated in the isolation medium, supplemented by (to reduce the risk of contamination) 2% of Penicilline/Streptomycine (Life Technologies) and 0.5% Amphotericin B (Sigma-Aldrich), at 37°C, 5% CO_2_. Medium was replaced after 24h to remove the non-attached cells and any other contaminant from the swab sample.

#### 2. Cell expansion

Expansion medium (PneumaCult Ex+ Medium, StemCell, with 1% of Penicilline/Streptomycine) was used for further culture of the cells and was changed 3 times a week. Once confluence was reached, cells were passaged using the animal component-free cell dissociation kit (StemCell). Cells were first incubated with the animal component-free (ACF) enzymatic dissociation solution (StemCell) on the cells for 15-20 minutes (min) at 37°C until they detached and then with ACF enzyme inhibiting solution (StemCell) to stop further dissociation. The cells were collected in a tube and centrifugated for 10 min at 1000 rotations per minute (rpm). The supernatant was discarded, and the cells were resuspended in a fresh expansion culture medium, seeded in a pre-coated flask, and incubated at 37°C until the cells became at least partly confluent. Meanwhile, the medium was changed every 2-3 days. Once confluent, using the same detachment protocol as above, 5×10^5^ cells in 200μL of expansion culture medium were seeded in the apical chamber of a transwell insert (24-well plate; 6.5 mm insert; 0.4 µm polyester membrane, Costar, Corning) while the basal chamber was supplemented by 700μL of the same medium. The inserts were then incubated at 37°C, the apical and basal media were changed 3 times per week. For preservation of the expanded cells to secure stocks, expanded cells were centrifugated for 10 min at 1000 rpm, resuspended in Recovery cell culture freezing medium (Thermofisher Scientific), and stored in liquid nitrogen. To use them, these cells were thawed and seeded in a medium flask with the expansion culture medium and later processed to transwells as described for non-frozen cells.

#### 3. *In vitro* cell differentiation

Once the cells reached confluency in the transwell, the expansion medium was removed from the apical and basal chambers. The differentiation process of the cells was started by culture at the air-liquid interface (ALI) at 37°C, where only the basal chamber was supplemented with 500μL of differentiation medium (PneumaCult ALI Basal Medium, StemCell, with 1% of Penicilline/Streptomycine). Medium was changed every 2-3 days. Full differentiation is reached after at least 3 weeks.

### *Ex vivo* culture of commercially available and in-house airway epithelia

Commercially available human airway epithelia (HAE) called “MucilAir™” purchased from Epithelix SARL [www.epithelix.com] were used as controls in some characterization experiments. According to the manufacturer and as previously described^17–19^, dedifferentiated epithelial cells obtained from nasal polyps were used to reconstitute *in vitro* these 3D tissues. Cells were cultured in air-liquid interface (ALI) at 37°C and 5% CO_2_ for 4-5 weeks, with 700μL of MucilAir™ medium (Epithelix) in the basal chamber. Once differentiated, the tissues comprise approximately 400’000 cells, of which around 200’000 are accessible from the apical surface. The MucilAir™ tissues are sTableduring months when cultivated in ALI at 37°C and 5% CO_2_. The availability of such tissues from children (contrarily to adults) is limited. For this reason, we decided to reconstitute our own HAE for this work. Once *in vitro* differentiated, the same *ex vivo* culture conditions were used to infect and maintain the in-house HAE.

### Respiratory virus tests

To confirm the absence of an asymptomatic respiratory viral infection during cell collection from donors (exclusion criteria), multiplex testing using semi-quantitative PCRs and RT-PCRs with a Euroscript kit (Eurogentec) was performed on primary swabs or freshly isolated cells for 15 viruses: Influenza A, Influenza B, RSV, Metapneumovirus, Picornaviruses (including Rhinovirus and Enterovirus), Para-influenza 1, 2, 3, 4, Adenovirus, Bocavirus and the four common cold Coronaviruses: HKU1, NL63, OC43, and 229E. The presence of SARS-CoV-2 was tested separately using the same protocol as virus quantification in the infection experiments (see below).

### Transepithelial electrical resistance (TEER) measurement

The TEER was measured at different timepoints during differentiation, as previously described^18^. Briefly, 200μL of differentiation medium was added in the apical chamber of the tissues. The resistance was measured with an EVOMX volt-ohmmeter (World Precision Instruments) and then converted into TEER using this formula: TEER (Ω.cm2) = (resistance value (Ω) - 100(Ω)) x 0.33 (cm2), where 100 Ω and 0.33 cm^2^ are the resistance and total surface of the membrane.

### Cilia beating assessment

Cilia beating of the reconstituted tissues was confirmed and recorded using a Sony XCD-U60CR microscope (Sony), Sony zcl software, and ImageJ software (National Institutes of Health), as previously described^18^.

### Immunofluorescence and hematoxylin and eosin staining

Tissues were fixed 30 min in paraformaldehyde 6% and then stored in PBS at 4°C. Staining was performed either directly on the whole tissue or on paraffin-embedded sections of 5μm for horizontal or transversal staining, respectively. For the immunofluorescence staining, tissues/sections were first permeabilized in Perm Wash buffer (BD Phospflow^TM^, BD Biosciences) 10 min at room temperature. Ciliated cells, goblet cells, basal cells, and tight junctions were stained using anti-β IV tubulin antibody (Ab179504, Abcam), anti-mucin 5AC (muc5AC) antibody (Ab3649, Abcam), anti-cytokeratin 5 (KRT-5) conjugated antibody (Ab193895, alexafluor 647; Abcam), and anti-zonula occludens-1 (ZO-I) antibody (33-910; Thermofisher scientific) respectively. Alexafluor anti-rabbit 488 antibody (A11032, Life Technologies), alexafluor anti-mouse 594 (A11032, Molecular Probe), and alexafluor anti-mouse 647 (A31571, Invitrogen) were used as secondary antibodies. Images were acquired with Zeiss LSM700 Meta Confocal Microscope (Zeiss) and analyzed by Imaris (Bitplane). Paraffin-embedded sections were also stained with hematoxylin (Merck) and eosin (Sigma).

### Gene expression of cell markers

Tissues were lysed during the differentiation (0 days=1week, 10 days=1.5week and 21 days=3weeks post-ALI transition). Total intracellular RNA was extracted (NucliSens easyMAG, BioMérieux) and the gene expression of KRT5, Forkhead Box J1 (FOXJ1), muc5A and neurogenic locus notch homolog protein 2 (Notch 2) was measured by semi-quantitative real-time RT-PCR using commercially available gene expression assay kits (Life Technology). Gene expression was normalized to Ribonuclease P expression (RNAse P) (Life Technology) as a housekeeping gene.

### Viruses

The viruses used in this study are summarized in Supplementary TableS2. IAV and RSV were isolated from clinical samples in commercially available HAE (MucilAir^TM^), and SARS-CoV-2 variant EG.5.1 in Vero-E6 cells overexpressing Transmembrane Protease Serine 2 (TMPRSS2) as previously described ^17,20^. RSV-A mCherry is a recombinant strain^21^. A HeLa cell-adapted clinical strain of RV A16 was purchased from ATCC.

### Infection

Infections of the fully differentiated nasal airway tissues were performed in ALI condition as previously described^17–19^. The multiplicity of infection (MOI) was defined according to the optimal kinetics of each virus, as in^17–19^ (TableS2). All MOIs were estimated based on an approximate number of accessible epithelial cells of ca. 200’000 per transwell^18^. Briefly, the tissues were transferred in fresh basal differentiation medium and incubated with PBS with calcium and magnesium 30 min at 37°C and 5% CO_2_ in order to strengthen the cell junction and thus prevent the basal migration of the virus. Then, after removing the PBS, the tissues were apically inoculated with 100μL of differentiation medium containing the virus. After 3 hours of apical virus adsorption at 33°C and 5% CO_2_, the tissues were washed 3 times with differentiation medium and incubated at 33°C and 5% CO_2_.

For the replication assessment at each timepoint, 200μL of differentiation medium was added apically and collected after 20 minutes of inoculation at 33°C and 5% CO_2_ and the basal medium was changed. For the assessment of host response by real time RT-PCR or RNA sequencing, virus- and mock-infected epithelial cells were lysed for each tested time point using easyMAG lysis buffer (BioMérieux).

### Assessment of virus replication and host response

Viral load was measured in the apical wash collected at each timepoint. RNA was extracted with NucliSens easyMAG (BioMérieux) and quantified by quantitative real-time reverse transcriptase polymerase chain reaction (RT-qPCR) using SuperScript™ III Platinum™ One-Step RT-qPCR Kit (Invitrogen) in CFX96 Thermal Cycler (BIORAD). RT-qPCR was performed using specific sets of primers and probes as previously described ^18,22,23^. Data were analyzed using Bio-Rad CFX maestro software (BIORAD). An apical wash similarly collected 3 hours post-infection (hpi) was considered the baseline for the assessment of viral replication.

Infected and non-infected tissues were lysed at 6hpi, 24hpi or 72hpi. Total intracellular RNA was extracted (NucliSens easyMAG, BioMérieux), and the induction of interferon (IFN) -α, -β and -λ and IFN stimulated gene 15 (ISG15) was measured by semi-quantitative real-time RT-PCR using commercially available gene expression assay kits (Life Technology). Gene induction in infected tissue is expressed in log10 fold change relative to non-infected tissue from the same donor and normalized to RNAse P as a housekeeping gene (Life Technology). Baseline levels of these genes in non-infected tissues were represented in cycle threshold normalized to RNAse P (ΔCT).

### Bulk RNA sequencing

RNA was extracted from cell lysates collected at different time points from mock- and infected HAE using TRIzol® Reagent (Invitrogen), according to the manufacturer’s instructions. Briefly, 720μL of TRIzol and 2μL of glycogen (ThermoFisher) were mixed with 120μL of the sample and incubated 5min at room temperature. Total RNA was quantified with Qbit (fluorimeter from Life Technologies) and RNA integrity was assessed with Bioanalyser (Agilent Technologies). Reverse transcription and cDNA amplification were performed with the SMART-Seq mRNA kit from Clontech, according to manufacturer’s specifications, starting with 1 ng of total RNA as input. Two hundred pg of cDNA were used for library preparation using the Nextera XT kit from Illumina. We assessed the library molarity and quality with the Quibit and Tapestation using a DNA High sensitivity chip (Agilent Technologies). Libraries were sequenced on a NovaSeq 6000 Illumina sequencer for SR100 reads.

### Transcriptomic analysis

Raw RNA-seq reads were processed with the nf-core rnaseq pipeline (3.14.0)^24^. Briefly, read quality control was performed with FastQC (0.12.1)^25^. Quality and adapter trimming was performed with Trim Galore (0.6.4)^26^. Read pseudoalignment and quantification were performed with Salmon (1.10.1)^27^ using the human genome GRCh38 (release 108 from Ensembl). Post-processing quality control reports were generated to assess the overall quality of the RNA-seq sample data.

Raw counts from protein coding genes were used to perform differential gene expression analysis with limma (3.58.1)^28^. Low-expression genes were removed with the filtered.data() function from the NOISeq R package (2.46.0)^29^. Raw counts were then normalized by calculating normalization factors with the calcNormFactors() function from the edgeR package (4.0.16)^30^ using the “TMM” method. Counts were transformed with the voom() function from the limma package prior to linear modelling. The subject from which samples were obtained was used as a random effect in the linear modelling step to account for inter-subject correlation when necessary. Moderated t-statistics, log2 fold-change, and Benjamini-Hochberg adjusted P values were obtained with the lmFit() and eBayes() functions. Significant differentially expressed genes were determined as those with an absolute log2 fold-change value > 1 and an adjusted P value < 0.05. Heatmaps of mean gene expression per group were generated with the ComplexHeatmap package (2.18.0)^30^. The list of genes selected according for their involvement in pathways related to IFNs and inflammation or their specific expression in ciliated, basal and goblet cells are summarized in Supplementary TableS3.

Gene set enrichment analyses were performed with the GSEA() function from the clusterProfiler package (4.10.1)^31^ using gene sets obtained with the msigdbr package (7.5.1)^32^. Significantly enriched terms were considered as those with an adjusted P value < 0.05.

From the same set of genes represented in the heatmaps (listed in Supplementary TableS3), the ones with a log2 fold-change value > 1 (upregulated) or < -1 (downregulated) were selected as input to perform over-representation analyses (ORA) with the clusterProfiler package (4.10.1)^33^.ORA results were obtained using the compareCluster() function by applying the enricher() function to a list of ORA inputs (gene IDs).

## Results

### HAE model of children and adults showed indistinguishable morphology and differentiation kinetics but differential expression of genes related to immunity and epithelial cell types

In total, fully differentiated HAEs were successfully reconstituted from nasal cells of 10 children (age median: 7 years, interquartile range: 6-9 years) and 9 adults (age median 38 years, interquartile range: 34-41 years) (Supplementary TableS1). The obtained tissues recapitulated the pseudostratified morphology and the cell composition of the *in vivo* nasal airway epithelium, as assessed by hematoxylin- and eosin-stained and immuno-stained slices of tissue (Fig.1A-B and Supplementary Fig.S1A). In these images, the 3 main cell types of the upper airway epithelium could be identified: ciliated, goblet, and basal (progenitor) cells. Cilia presence and beating at the HAE apical surface were also confirmed in the recorded movie from fully differentiated tissues (supplementary material). Further characterization of tissues reconstituted from adults and children included a time-course study of the tissue integrity and the expression of differentiated cell markers at the beginning, during, and at the end of the differentiation, respectively at week(s) 0, 1.5, and 3, post-ALI transition (wpa). As shown in Figure 1C, increasing TEER was observed and all tissues reached the integrity threshold (commonly^18,34^ fixed at 100 Ω.cm^2^) between 0 and 1.5 wpa, despite inter-donor variability. Except for one from an adult donor with higher TEER, all tissues displayed comparable TEER between children and adults, ranging from 151.8 to 839.5 Ω.cm^2^ at 1.5wpa and from 123.8 to 740.2 Ω.cm^2^ at 3wpa. This result, in addition to the comparable expression levels of RNAse P gene used as an internal control (Supplementary Fig.S1B), indicate no major difference in the thickness of these epithelial tissues. Tissue integrity was also confirmed through the presence of tight junctions by immuno-staining using an anti-ZO-I antibody (Fig.1D-E) at 3wpa. Automatic reconstitution of the tissue from immunofluorescence images highlighted the progressive increase of tissue thickness during the differentiation (Fig.1F), with distribution evolving from monolayer at 0wpa to pseudostratified morphology at 1.5 and 3wpa. At 0wpi, only basal cells (expressing KRT5 marker) were present in HAE from children and adults, contrary to 1.5 and 3wpa, where a gradual increase in ciliated and goblet cell markers (respectively β IV tubulin and muc5AC markers) was detected. Expression of epithelial cell type markers and four IFN related genes was assessed by RT-PCR (Fig.1G-I and Supplementary Fig.S1C-G). Although not statistically significant, we observed the trend of higher expression of only KRT5, FOXJ1, and muc5AC, targeting KRT5, FOXJ1 and muc5AC, for basal, ciliated, and goblet cells, respectively^35,36^. The expression of these three markers seems to peak at 1.5wpa and then slightly decreases at 3wpa.

**Figure 1:**
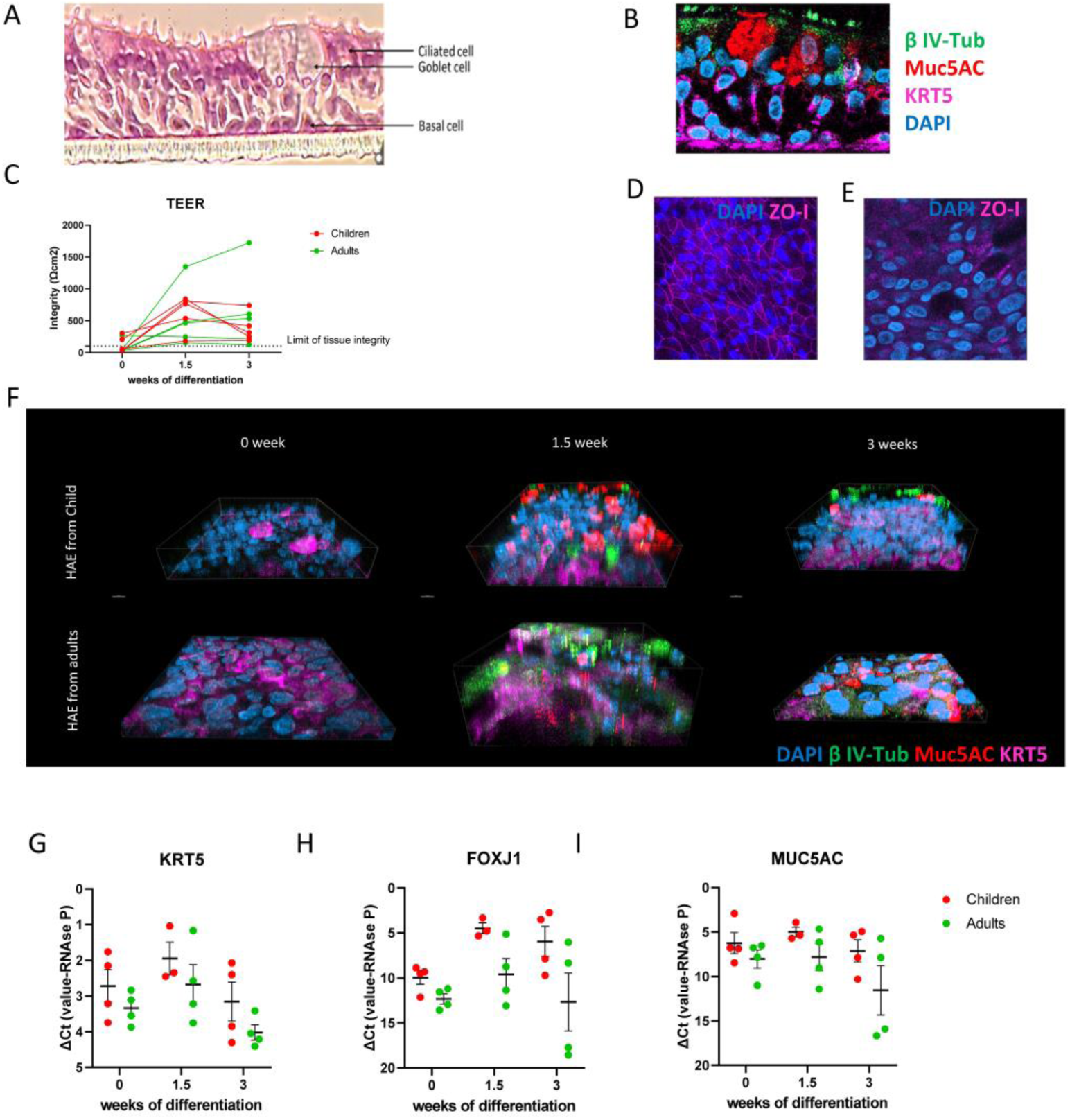
Morphological and functional characterization of the model in the differentiation course. Isolated nasal epithelial cells were cultured and used for HAE *in vitro* differentiation. **A:** Hematoxylin- and eosin-stained transversal section of a fully differentiated child tissue with the three different cell types composing the nasal epithelium. **B:** Section of the same fully differentiated child tissue stained by immunofluorescence. Blue: DAPI (nuclei), green: β IV-tubulin (β IV-Tub, ciliated cells), red: mucin 5AC (muc5AC, goblet cells), pink: keratin 5 (KRT5, basal cells). **C:** Transepithelial electrical resistance (TEER) measurement in tissues at 3 different stages of differentiation (0, 1.5, and 3 weeks post-ALI transition (wpa)). N children=5 N and adults=5. **D, E:** Immunofluorescence of fully differentiated adult (D) and child (E) tissues. Blue: DAPI (nuclei), pink: zonula occludens-1 (ZO-I, tight junction). **F:** Immunofluorescence of a child (upper panels) and an adult (lower panels) tissues at 3 different stages of differentiation: 0 (left panels), 1.5 (middle panels), and 3 (right panels) wpa. Blue: DAPI (nuclei), green: β IV-Tub (ciliated cells), red: muc5AC (goblet cells), pink: KRT5 (basal cells). Images were acquired with Zeiss LSM700 Meta Confocal Microscope and used for 3D reconstitution by Imaris (V9.6) software. **G, H, I**: Gene expression of KRT5 (expressed by the basal cells, panel G), Forkhead Box J1 (FoXJ1, expressed by the ciliated cells, panel H) and muc5AC (expressed by the goblet cells, panel I) in tissues from children (N=4) and adults (N=4) at different stages of differentiation (0, 1.5, and 3 wpa) measured by RT-PCR. All the values were normalized with the expression level of RNAseP (internal control). The mean and standard error of the mean (SEM) are represented by black lines. No statistically significant difference was found.

In line with these observations suggesting that most of HAE differentiation, from both child and adult nasal cells, occurs during the first 10 days (between 0 and 1.5wpa). Transcriptomic analysis performed on HAE from children and adults at the same time points showed that, in both age groups, many genes were differentially expressed at 0wpa compared to 1.5wpa and 3wpa. In contrast, no statistically significant change in terms of gene expression was observed between 1.5wpa and 3wpa in HAE from both age groups (Supplementary Fig.S2A-F). Comparing the HAE transcriptomic pattern in children versus adults at the three differentiation stages (0, 1.5 and 3wpa), principal components analysis (PCA in Supplementary Fig.S2G) showed no clear clustering between the two age groups. A better separation between children and adults was observed for most donors at 1.5 and 3wpa compared to 0wpa. This was likely associated with wider differences in pathway induction at these two timepoints compared to 0wpa, where only subtle changes were observed between pediatric and adult donors (Fig.2A and Supplementary Fig.S2H-I). In particular, a progressive enrichment in adults (negative normalized enhancement score in the x-axis) of pathways related to interferon response and inflammation was observed from 1.5wpa and was even more significant at 3wpa. In line with these observations, an overall higher expression of genes related to IFN pathways and inflammation was found in adults HAE at 1.5 and 3wpa (Fig.2B-C and Supplementary Fig.S3). Upregulation of genes specific to ciliated and goblet cells was observed in children while genes specific to basal cells were upregulated in adults (Fig.2D-F and Supplementary Fig.S3). Looking at host factors involved in virus entry (Fig.2G), we found no significant difference in the expression of ACE-2 (angiotensin-converting enzyme 2) and TMPRSS2, involved in SARS-CoV-2 entry, ICAM-1 (InterCellular Adhesion Molecule 1) for RV, and CX3CR1 (C-X3-C Motif Chemokine Receptor 1), an RSV receptor, between children and adults.

**Figure 2:**
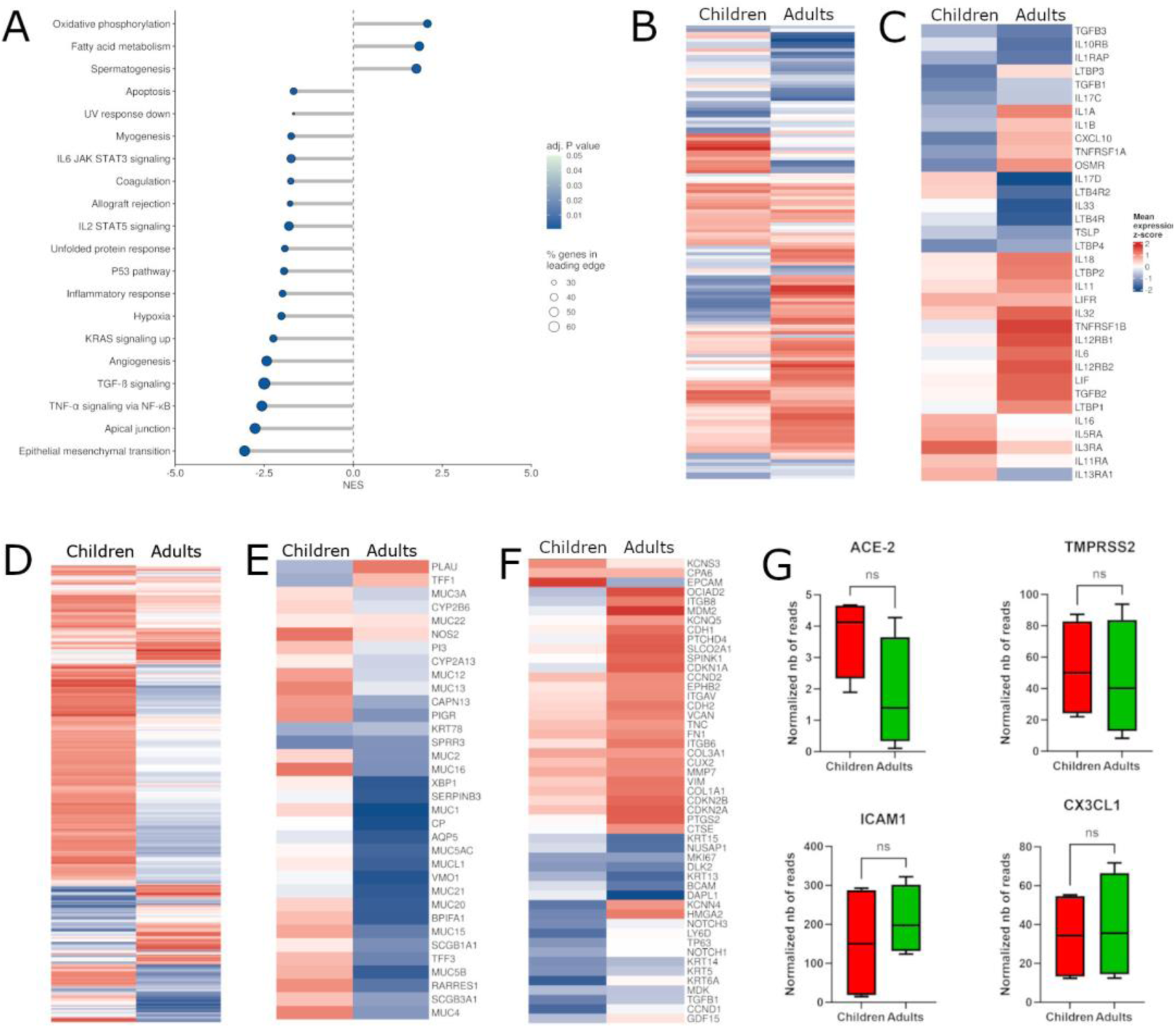
Differential pathways induction in HAE from children versus adults at the end of the *in vitro* differentiation. **A:** Hallmark gene set enrichment analysis (GSEA) of fully differentiated tissues from child compared to adult at 3wpa. Positive normalized enrichment score (NES): pathways upregulated in children, negative NES: pathways upregulated in adults. Size of the dots: percentage of genes in the leading edge (from 10 to 40%). Color of the dots: adjusted P value (from 0.01 (darkest blue) to 0.05 (lightest blue)). **B, C, D, E, and F:** Heatmaps of a subset of selected genes of interest related to interferon response (B), inflammation (C), ciliated cells (D), goblet cells (E) and basal cells (F) at 3wpa. Color: mean expression z-score. **G:** Expression (normalized number of reads) of host factors involved in entry, ACE-2 and TMPRSS2 (for SARS-CoV-2), ICAM1 (for RV), and CX3CR1 (for RSV). In all panels, N adults=4, N children=4. See Supplementary Fig.S3 for more details

Altogether, the characterization of HAE reconstitution from children and adult nasal epithelial cells revealed similar differentiation kinetics and no major visible morphological differences between the two age groups of donors. However, during and at the end of the differentiation, an increasing induction level of pathways potentially involved in the mucociliary clearance (MCC) function was observed in epithelial cells from pediatric donors, while higher upregulation of innate immune response was found in HAE from adult donors.

### Transcriptomic analysis of respiratory infections in nasal HAE from adults and children revealed differential host gene induction, despite comparable viral replication between the two age groups

Next, we studied age-related gene induction in fully differentiated HAE in the context of respiratory viral infection with different viruses. To validate the HAE model as a tool to compare respiratory viral infections *ex vivo*, we first confirmed comparable infections by infection with RSV in our in-house HAE, compared to commercially available ones, both from adult donors (Supplementary Fig.S4A-B). Then, infection conditions and approach (Supplementary Fig.S4C-F) were tested using RV and IAV before further assays using more viruses. In these preliminary tests in our in-house HAE from children versus adults, gene expression assessed by RT-PCR (Supplementary Fig.S4G-N) already suggested slight differences between the age groups regarding host response, an assumption that needed further confirmation and deeper investigation by unbiased methods and involving more donors.

Giving these first observations, virus infection assays in HAE from children and adults with IAV, RSV, RV and SARS-CoV-2 were conducted under the same conditions (Supplementary Fig.S4C-D), but here analyzed using a non-focused approach with transcriptomic analysis. Information about viral strains is summarized in Supplementary TableS2. Virus-specific MOIs were selected according to previous studies to have the optimal infection conditions^17–20^ and to assess host response during the exponential phase of virus replication. In addition, replication kinetics for each virus were measured in separate epithelial tissues of the same donors in parallel to HAE infections for the transcriptomic analysis (Supplementary Fig.S5A-D). RV and IAV replicated faster than SARS-CoV-2 and RSV, with a maximal replication at 48hpi versus 72hpi. During the peaks of infections, the levels of replication were higher for IAV (8.3 log10 RNAc/mL for children and 8.4 log10 RNAc/mL for adults) and SARS-CoV-2 (9.0 log10 RNAc/mL for children and 10.2 log10 RNAc/mL for adults), compared to RV (7 log10 RNAc/mL for children and 6.9 log10 RNAc/mL for adults) and RSV (5.8 log10 RNAc/mL for children and 4.9 log10 RNAc/mL for adults). Nevertheless, viral replication shows similar kinetics in HAE from children compared to adults for all tested viruses. Viral replication was also checked in each tissue used for the transcriptomic analysis and showed no significant difference in the levels of viral RNA (Supplementary Fig.S5E-H) and infectious titer (Supplementary Fig.S5I-L) between children and adults for each virus.

To consider the stress exerted on HAE response during its manipulation (inoculation, washes, and virus collection at each time point) and that may provoke a host response independent of the virus infection, a mock-infected tissue (using a virus-free medium) was compared to virus-infected tissue at each time point. Despite variations between 6, 24, and 72hpi, most probably due to this stress (Supplementary Fig.S5I-K), the overall pathways enrichment pattern between children and adults was comparable to what was previously observed during tissue differentiation (Fig.2). After gene expression normalization to mock infection (to focus on the host response specifically induced during viral infection), we next performed gene set enrichment analysis (GSEA) to identify pathways differentially regulated between the two age groups (Fig.3). From the first hours of infection by SARS-CoV-2, several pathways related to immune response showed higher activation in children (normalized enrichment score higher than 0 in the x-axis). The highest enrichment was observed with type I and II IFN responses throughout all three timepoints (Fig.3A-C). In contrast, genes related to complement and interleukin-6 JAK-STAT (Janus Kinase-Signal Transducers and Activators of Transcription) signalling were only enriched in children at 6hpi and genes related to apoptosis, NF-κB (nuclear factor-κB)-mediated TNF-α (Tumour Necrosis Factor α) signalling, inflammatory response and TGF-β (transforming growth factor β) signalling until 24hpi (Fig.3A and B). SARS-CoV-2 showed the most pronounced and earliest age-related divergence in pathways induction compared to RV, RSV, and IAV, where, except for IFN-α in RV- and IAV-infected HAE, the immune response was higher in adults at 6hpi (Fig.3D-L). Although observed later, some of the pathways enriched in HAE from children infected with SARS-CoV-2 were also found in the case of HAE RV infection but only at 24 and/or 72hpi (Fig.3E and F). This includes IL-6 JAK-STAT3, NF-κB-mediated TNF-α, TGF-β signalling, and inflammatory response. However, in the context of RSV and IAV infections, most of these pathways involved in immune response were not enriched in children, except for TNF-α signalling at 72h post-RSV infection and 24h post-IAV infection, and TGF-β signalling at 24h post-IAV infection. Some pathways remained even higher in adults at 6hpi (like in the absence of infection). At 72hpi, IFN-α response was enriched in RSV-infected HAE from adults (Fig.3I), but in IAV-infected HAE from children (Fig.3L). Pathways related to IFN-γ response also showed higher induction in children only at 72h post-IAV infection (Fig.3L).

**Figure 3:**
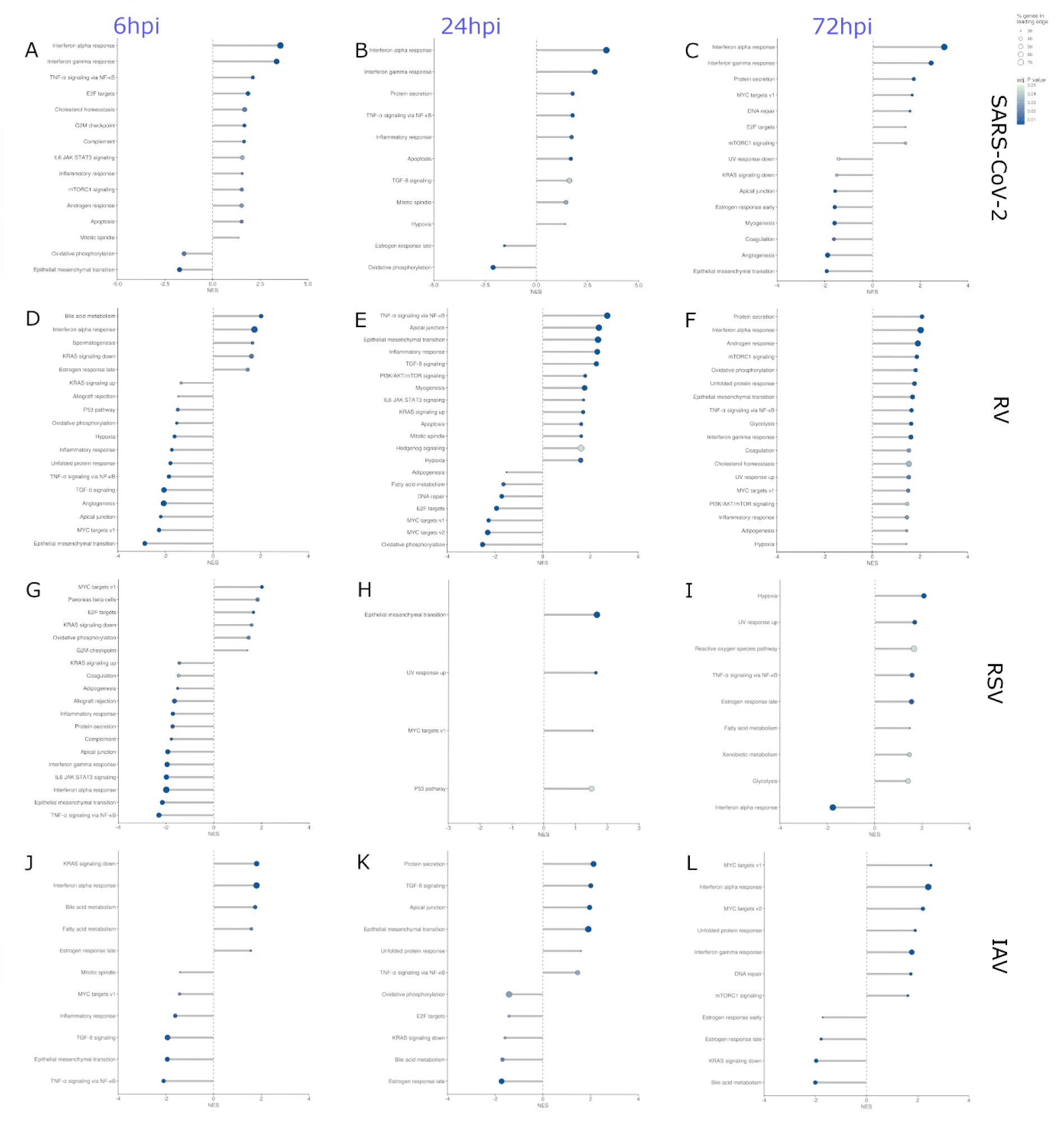
Differential pathways induction in HAE from children versus adults infected by SARS-CoV-2, RV, RSV and IAV. **A, B, C:** Hallmark gene set enrichment analysis (GSEA) of SARS-CoV-2 infected child tissues (N=5) compared to SARS-CoV-2 infected adult tissues (N=3) at 6 hours post-infection (hpi) (A), 24hpi (B), and 72hpi (C). **D, E, F:** Hallmark GSEA of RV-infected child tissues (N=5) compared to RV-infected adult tissues (N=4) at 6hpi (D), 24hpi (E), and 72hpi (F). **G, H, I:** Hallmark GSEA of RSV-infected child tissues (N=3) compared to RSV-infected adult tissues (N=3) at 6hpi (G), 24hpi (H), and 72hpi (I). **J, K, L:** Hallmark GSEA of IAV-infected child tissues (N=7) compared to IAV-infected adult tissues (N=5) at 6hpi (J), 24hpi (K), and 72hpi (L). The gene expression of each donor was normalized with a mock-infected tissue from the same donor, collected at the same timepoint. Positive normalized enrichment score (NES): pathways upregulated in children, negative NES: pathways upregulated in adults. Size of the dots: percentage of genes in the leading edge (from 10 to 40%). Color of the dots: adjusted P value (from 0.01 (darkest blue) to 0.05 (lightest blue)).

To complement the GSEA results, we assessed the expression (normalized to non-infected HAE from each donor) of genes related to each epithelial cell type or involved in inflammation and interferon response, using the same samples. As shown in Figure 4A (and Supplementary Fig.S6 with more details), an early and strong induction of genes involved in IFN response was specifically found in SARS-CoV-2 infected HAE from children from 6 to 72hpi. This includes genes encoding sensors like IFIH1 (for MDA-5, melanoma differentiation-associated protein 5) and DDX58 (for RIGI, retinoic acid-inducible gene I), proteins involved in signalling downstream and upstream IFN activation like IRF1, 3 and 9 (IFN-regulatory factors) and STAT1, 2 and 3 (signal transducer and activator of transcriptions), and IFN-stimulated genes like ISG15, MX1 (Myxoma Resistance Protein 1) and 2, OAS (2′,5′-oligoadenylate synthetase) and chemokines like CXCL11 (chemokine C-X-C motif ligand 11). A slighter induction of some of these genes was also observed in child tissues infected by IAV and RV, with a progressive increase until 72hpi with IAV.

**Figure 4:**
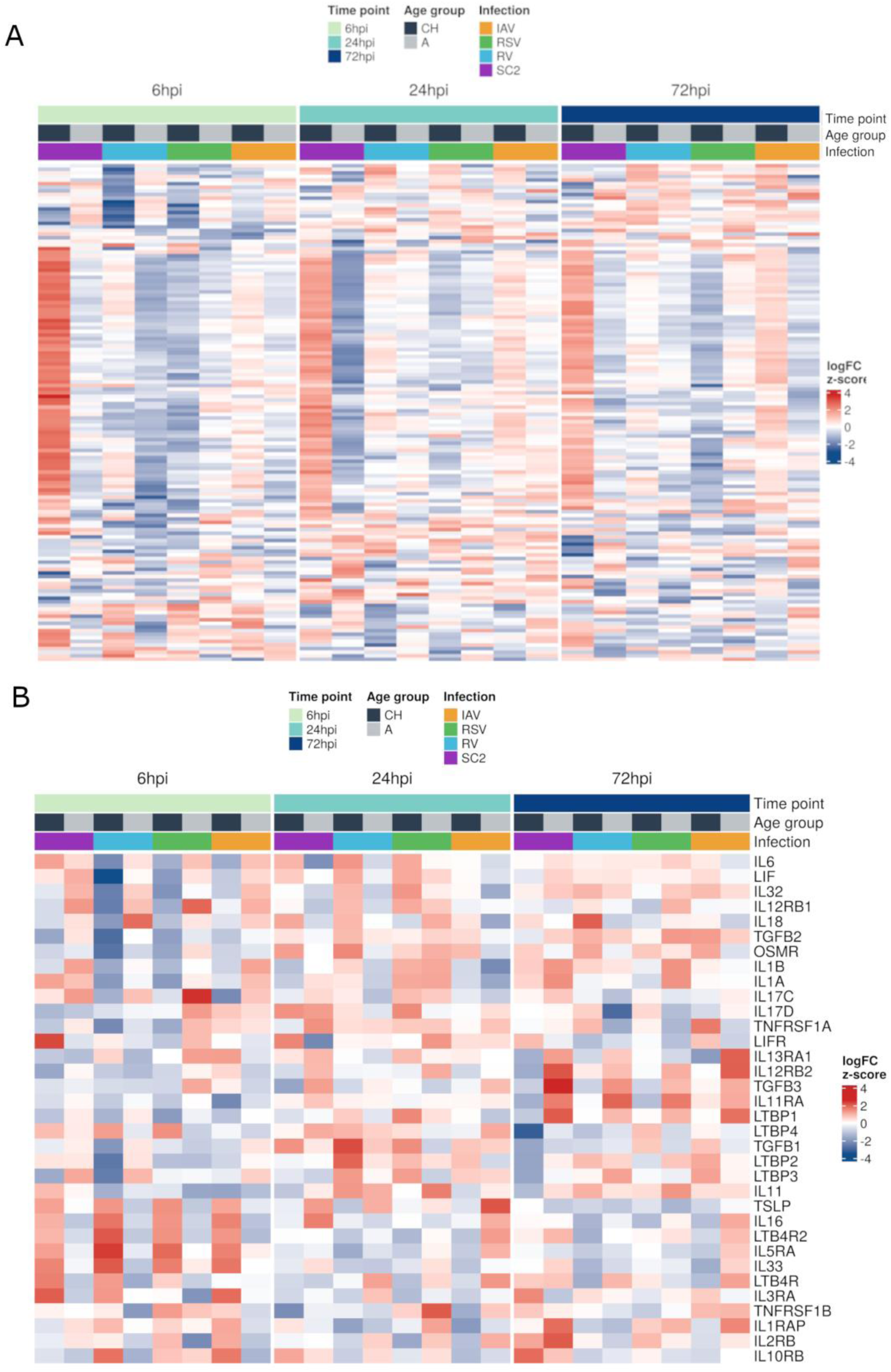
Differential expression of genes involved in immune response in HAE from children versus adults infected by SARS-CoV-2, RV, RSV and IAV at 6hpi, 24hpi, and 72hpi: Heatmaps of selected genes involved in interferon signalling pathways (A) and in inflammatory response (B) in tissues infected by SARS-CoV-2, RV, RSV, and IAV, normalized with the gene expression in the corresponding mock-infected tissue from the same donor and at the same time point, in children and adults (N: similar as in figure 3 for the different viruses) at 6hpi, 24hpi, 72hpi. See Supplementary Fig.S6 for more details

Among genes involved in inflammation (Fig.4B), interleukin 6 (IL-6) was more expressed during the first hours of infection (6hpi) in SARS-CoV-2 infected child tissues, unlike the other viruses. Conversely, some genes like IL-33, IL-11, IL-16 and TSLP (Thymic stromal lymphopoietin) showed higher induction in children by all respiratory viruses only at 6hpi. Their induction pattern between children and adults is even inverted at later timepoints in IAV-infected cells. Induction of ILs -32, -18, -1α, -1β, -17C and - 17D were higher in HAE from adults infected by IAV and SARS-CoV-2 at 6hpi. At 24hpi, these genes were mainly induced by RV in children and by RSV in both age groups (except IL-18 in children). TGF-β3 (Transforming growth factor beta 3), LTBP-1 (latent transforming growth factor-β–binding protein) and some IL receptors IL13RA1, IL12RB2, IL11RA showed higher induction in HAE from adults compared to children with all studied viruses at 72hpi. For these genes, the most and least pronounced differences between the two age groups were observed, respectively, in SARS-CoV-2- and IAV-infected HAE at 72hpi.

We also assessed the expression of genes specifically expressed by the different epithelial cell types (Supplementary Fig.S7). Higher expression of genes involved in the MCC in children observed at the end of the differentiation was also observed at 6hpi with all viruses, especially in case of RV infection, but less with RSV where more induction of genes related to mucus production in adults at 6 and 24hpi was observed. Upregulation of genes related to goblet cells in adult donors was also observed at 72hpi in the case of SARS-CoV-2 and IAV infections.

Finally, using the same set of genes related to IFN pathways and inflammation where we observed differences between the two age groups in infected HAE (used in Fig.4 and S7 and listed in supplementary Supplementary TableS3), we determined the related pathways that were specifically up- or down-regulated in children compared to adults at each time point (Fig.5). At a glance, we could confirm the overall distinct pattern SARS-CoV-2, compared to the other respiratory viruses and especially RSV and IAV in infected HAE, mostly with pathways related to immune response. This difference was again more pronounced at the earliest infection time point, 6hpi. In particular, pathways related to inflammation and TNF-α signalling via NF-KB were upregulated in case of SARS-CoV-2 infection but rather down-regulated for RV, RSV and IAV in children at 6hpi.

**Figure 5:**
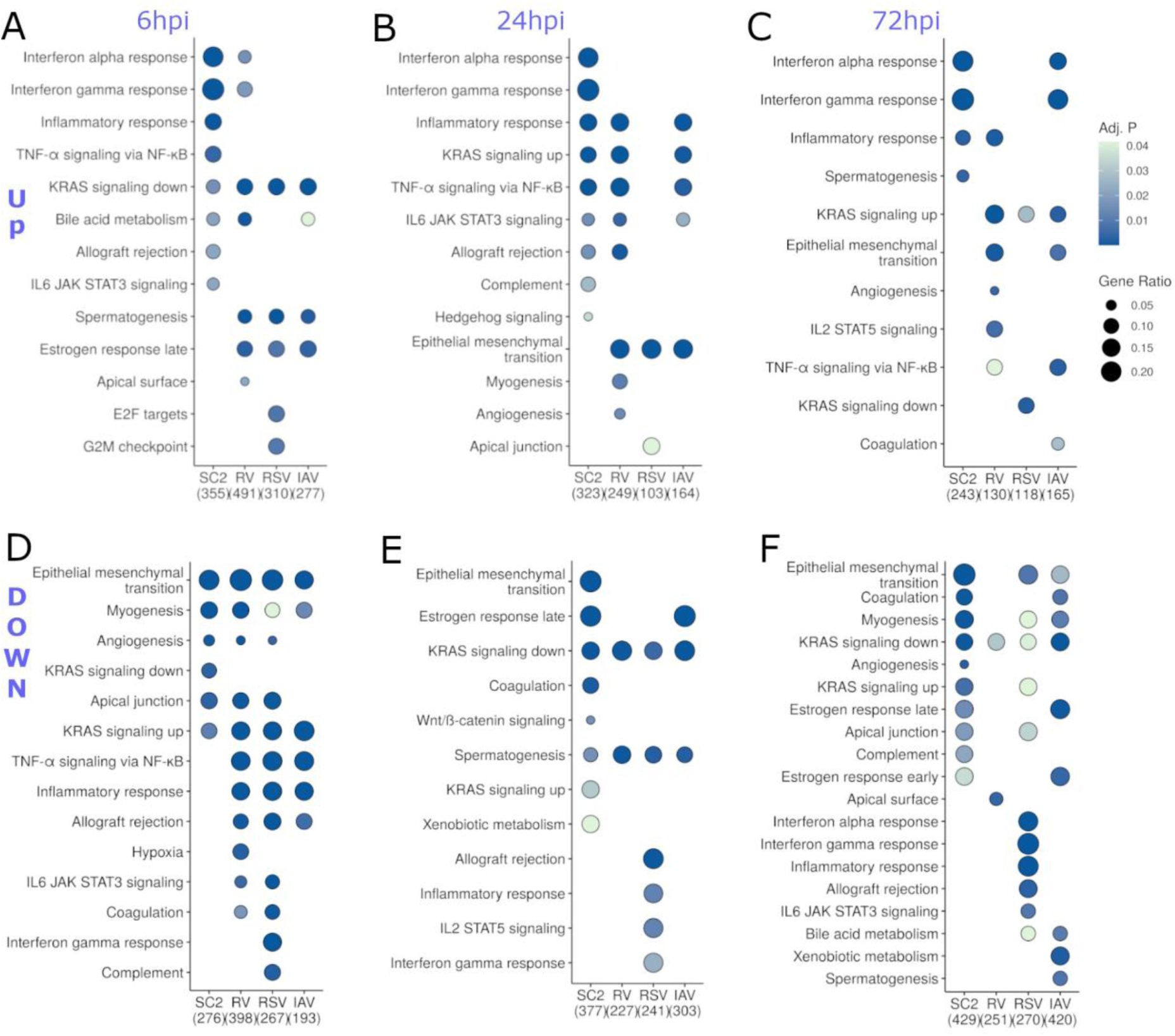
Enriched pathways involved in immune response in HAE from children versus adults infected by SARS-CoV-2, RV, RSV and IAV at 6hpi, 24hpi, and 72hpi: Hallmark over-representation analysis (ORA) of pathways up (A, B, and C) or downregulated (D, E, and F) in child compared to adult tissues in the context of infection by SARS-CoV-2 (SC2), RV, RSV, and IAV at 6hpi (A and D), 24hpi (B and E), and 72hpi (C and F). Normalization with the gene expression in the corresponding mock tissue. N: as in figure 3 for the different viruses).

To conclude this part, despite similar viral replication of all respiratory viruses between the two age groups, MCC functions and immune response implicated in anti-viral state and/or inflammation induction appeared differentially regulated in children compared to adults and depend on the timing of the infection and the infecting virus species.

## Discussion

The pathogenesis of respiratory viruses is tightly linked to their interaction with the human airway epithelium during the early stage of infection. The related disease outcomes rely not only on the virus species but also on host responses, which vary over lifetime or become modulated in the context of comorbidity^37,38^. This work represents one of the few studies investigating four highly prevalent and clinically relevant respiratory viruses in an *in vitro* model of the nasal epithelium of children and adults. In addition to the comparison by age, it includes a comparison of HAE response to four of the most clinically relevant respiratory viruses.

Nasal human airway epithelia constitute one of the primary sites of respiratory viral infections, with the respiratory epithelium comprising the first line of defence. In addition to the physical barrier insured by the MCC function, the recognition of viruses through pattern recognition receptors (PRRs) like Toll-like receptors (TLRs) triggers signalling pathways that lead to the production of a plethora of anti-viral and proinflammatory mediators, including cytokines and chemokines, which recruit and activate immune cells involved in both innate and adaptive immunity (reviewed in ^16^). *In vitro* reconstituted HAE, cultured in air-liquid interface, is nowadays recognized as the “universal”^39^ model to study viral infection *ex vivo*, offering a closer approximation to human physiological conditions and bridging the gap between *in vitro* studies and animal models^39–42^. After the successful set-up and characterization of our in-house 3D culture model, our first comparison showed that, despite similar differentiation kinetics between children and adults, disparate induction levels of genes involved in the MCC and immune response between the two age groups were reached after the full tissue reconstitution between the two age groups. Previous analyses from *in vitro* and *in vivo* data showed different cellular compositions of the nasal epithelium between age groups, with some studies observing a higher number of goblet cells and a lower number of ciliated cells in children^15,43^, and others reporting a higher proportion of ciliated and goblet cells but less basal cells in the child nasal mucosa compared to adults^13,44^. Although not enough accurate for quantification, analysis by immunostaining of reconstituted HAE did not show marked differences in ciliated or goblet cell proportions between age groups in our model. However, our transcriptomic analysis reported a greater induction of genes related to these two epithelial cell types in pediatric tissues, possibly associated with MCC function. Of note, MCC function was reported to be affected by respiratory viral infection^18,45,46^. Characterization of the single-cell transcriptomic landscape from nasal swabs collected on patients reported a pre-activation of the innate immune system before infection in non-infected children^13,15^. By RNA sequencing of tracheobronchial cells from children and adults, *Maughan et al.* recently showed that, in contrast to freshly sorted basal cells (*in vivo*), proliferating cultured basal cells (*in vitro*) showed more age-related transcriptional differences, including higher induction of genes related to TNFα signalling and the inflammatory response in adults, while these gene sets were slightly more expressed in freshly sorted pediatric basal cells (*in vivo*)^47^. Discrepancies between these *in vivo* versus *in vitro*/*ex vivo* data could be due to constant but reversible epithelial pre-activation of epithelial cells in children (*in vivo).* This phenotype might be driven by frequent prior stimuli, such as viral/bacterial infections and seems to be maintained even after pathogen clearance^48^. These discrepancies between the *ex vivo* and *in vivo* model could also be due to the absence of resident immune cells in cultured HAE that might as well play a role in the age-dependent immune response pre-activation, as recently suggested^49^. These data stress one limitation of our model in which no resident immune cells are present, thus while our study can elucidate the innate immune response on the epithelial level, the component of the cross-talk between epithelium and immune cells that is lacking.

While a large range of data were generated on SARS-CoV-2-host interactions during the COVID-19 pandemic, much less is known about this interplay in the context of other respiratory viruses, for which findings on SARS-CoV-2 cannot be necessarily extrapolated. Our data showed virus-specific HAE immune responses of the infected HAE model. From a clinical point of view, unlike Influenza Virus and RSV, SARS-CoV-2, on average, has a lower morbidity in children than in adults. The more pronounced clinical disease of RSV and Influenza Virus in children could also be reflected by higher and less and/or later control induction of inflammatory pathways and/or less efficient early innate immune responses. We showed heightened gene expression of pathways related to the IFN response in SARS-CoV-2 HAE in children but more pronounced upregulation of inflammation-related genes in HAEs from adults at later time points. Thes IFN pathways involve genes encoding for IFN sensors, IFN signalling and Interferon Stimulated Genes (ISGs). Compared to IAV, RV, and RSV, SARS-CoV-2 also showed the highest and earliest induction of innate immune response in children. Several hypotheses have been proposed to explain this age-related difference in terms of the clinical severity of SARS-CoV-2 infection. Some of them, such as the proposed lower level of viral replication or viral entry factors’ expression (namely ACE-2 and TMPRSS2), or a cross-protection due to previous infection by seasonal common cold coronaviruses in children, have been contradicted^50^. More recently, a number of studies using clinical samples from infected patients were rather supporting the importance of the interplay of the immune response in the modulation of COVID-19 disease severity in children compared to adults^14,51,52^. In particular, a more robust local innate immune response in children has been observed in nasal mucosa. Transcriptomic analysis of nasal swabs showed a higher expression of viral sensors like MDA-5 and RIG-I and of ISGs before infection and in the early phase of infection in children compared to adults. The presence of immune cells (especially neutrophils) was also higher in children’s nasal mucosa before infection^15,53^. At the level of the systemic immune response, children appear to better contain the inflammation and limit the disease course with a faster resolution of the antiviral state, a more naive repertoire of immune cells that are more transiently activated monocytes and dendritic cells, a more limited activation of the cytotoxic response and an earlier activation of the B cells^13,14^. Coherent with our findings, all these studies suggested that more potent activation of the nasal mucosal immune system in children is likely responsible for a faster but stronger response and better control of SARS-CoV-2 infection. Of note, age-dependent HAE response can vary across SARS-CoV-2 variants^54^. Using a more recent SARS-CoV-2 subvariant, EG.5.1, from the Omicron clade, our findings regarding the induction of genes related to IFN pathways, including sensors, proteins involved in signalling and ISGs globally confirmed previous investigations with earlier SARS-CoV-2 strains comparing infection in children versus adults^13,15^.

One of the rare studies comparing host response in children versus adults between respiratory viruses reported that SARS-CoV-2 was associated with only minimal viral sensing by bronchial HAE, compared to Influenza^55^. Using nasal HAE, we showed a rather disparate pattern provoked by SARS-CoV-2, especially compared to RSV and IAV. A comparison of the immunological features from nasopharyngeal swabs from subjects with Influenza versus RSV infections demonstrated severe disease development dependency on the host response rather than the virus nature^56^. An analysis of cytokines’ secretion by PBMC showed significantly different ratios of type 1 and 2 immune response modulators between RSV-infected children with URT infections versus bronchiolitis, leading to an excessive type 2/deficient type 1 cytokine response. This suggested an association of inappropriate cross-regulation between the two adaptive responses with enhanced severe infantile RSV infection in the respiratory tract ^11^. Similar conclusions have also been drawn from a study of lung tissues from fatal RSV and influenza LRT infection cases^9^. As RSV is more pathogenic in infants and the elderly, a similar investigation using older adults and younger children would be of interest.

Comparing the induction of genes and pathways in children to adults, our data showed that IAV and RSV had the least marked increase of immune response in pediatric donors during the three first days of infection. IAV also showed the most disparate profile of inter-age-group change in the expression of genes related to MCC and inflammation. RV showed an intermediate pattern closer to SARS-CoV-2 than RSV and IAV. *Usemann et al.* recently suggested that a slight increase of RV replication in *in vitro* reconstituted pediatric HAE could be related to a lower immune response at 20hpi^57^. Although no change was observed in terms of viral replication from 6 to 72hpi, we here reported that, after the early decrease during the first hours of infection, this response was progressively increased later compared to adults. RV mainly causes mild disease in children, where exacerbations have been described to be more associated with a history of respiratory disorders like allergy, cystic fibrosis, and asthma^58–60^. Although more frequent in children, RV clinically shows less age-dependent differences in terms of illness severity. On the contrary, adults were more associated with enhanced symptom severity and higher shedding was reported^61,62^. For all the studied respiratory viruses, controversial conclusions have indeed been reported regarding viral shedding in children versus adults ^22,63–70^. Viral shedding and viral load detection can be influenced by many other factors, like the clinical severity or underlying immunity, as well as the site of sampling. Despite the age-specific differential induction of host response, we here found that virus replication was comparable in HAE from children versus adults in the *ex vivo* model. The recent studies of HAE response to viral infection, including ours, suggest that differential virus-host interaction, rather than virus entry and replication, might impact clinical manifestation between the age groups.

In summary, our findings suggested that, while innate immune response seems more pre-activated in adults *ex vivo* before any prior infection, a virus-specific enhancement of host induction in children during the early stage of infection might drive a better control against SARS-CoV-2 than IAV, RSV and RV. This age-related signature seems to be more poised to better control the fate of the infection and could explain the mitigated clinical course in SARS-CoV-2 infection in children. Thus, our work supports conclusions regarding the age dependency of virus sensing and additionally demonstrates a distinct HAE stimulation by SARS-CoV-2 compared to pre-pandemic respiratory viruses. It furthermore provides evidence about the crucial role of the early viral interaction with the human airway to intrinsically mediate the activation of diverse immune response actors for even later stages of the infection. Beyond this project, this emphasizes again the relevance of such an *ex vivo* model as a powerful tool to mimic respiratory infections in patients and study viral pathogenesis.

## Acknowledgement

We thank the iGE3 genomics platform of the UNIGE (University of Geneva) for the sample sequencing, the Histology platform of the UNIGE for the preparation and colouration of the sliced tissues, and the Laboratory of Virology of Geneva University Hospitals for the clinical samples used for virus isolation. We also acknowledge Caroline Tapparel-Vu (UNIGE) for providing the RV ATCC strain, Meriem Bekliz (UNIGE) and Kenneth Adea (UNIGE) for the isolation of the SARS-CoV-2 EG5.1 strain, and Jean-Francois Eleouet (VIM UR892, INRAe, Université Paris-Saclay) and Marie-Anne Rameix-Welti (Université Paris-Saclay, INSERM, UVSQ, UMR 1173 (2I)) for the RSV-A mCherry virus. Finally, we thank all the child and adult donors for voluntarily participating in the study.

## Reference

1. Jin, X., et al. Global burden of upper respiratory infections in 204 countries and territories, from 1990 to 2019. eClinicalMedicine 37, (2021).

2. Galanti, M. et al. Longitudinal active sampling for respiratory viral infections across age groups. Influenza Other Respir Viruses 13, 226–232 (2019).

3. Sarna, M. et al. Viruses causing lower respiratory symptoms in young children: findings from the ORChID birth cohort. Thorax 73, 969–979 (2018).

4. Teoh, Z. et al. Burden of Respiratory Viruses in Children Less Than 2 Years Old in a Community-based Longitudinal US Birth Cohort. Clinical Infectious Diseases 77, 901–909 (2023).

5. Nair, H. et al. Global burden of acute lower respiratory infections due to respiratory syncytial virus in young children: a systematic review and meta-analysis. Lancet 375, 1545–1555 (2010).

6. Fraaij, P. L. A. & Heikkinen, T. Seasonal influenza: the burden of disease in children. Vaccine 29, 7524–7528 (2011).

7. Wang, X. et al. Global burden of respiratory infections associated with seasonal influenza in children under 5 years in 2018: a systematic review and modelling study. The Lancet Global Health 8, e497–e510 (2020).

8. Götzinger, F. et al. COVID-19 in children and adolescents in Europe: a multinational, multicentre cohort study. Lancet Child Adolesc Health 4, 653–661 (2020).

9. Welliver, T. P., Reed, J. L. & Welliver, R. C. Respiratory syncytial virus and influenza virus infections: observations from tissues of fatal infant cases. Pediatr Infect Dis J 27, S92–96 (2008).

10. Larrañaga, C. L. et al. Impaired immune response in severe human lower tract respiratory infection by respiratory syncytial virus. Pediatr Infect Dis J 28, 867–873 (2009).

11. Legg, J. P., Hussain, I. R., Warner, J. A., Johnston, S. L. & Warner, J. O. Type 1 and type 2 cytokine imbalance in acute respiratory syncytial virus bronchiolitis. Am J Respir Crit Care Med 168, 633–639 (2003).

12. Welliver, T. P. et al. Severe human lower respiratory tract illness caused by respiratory syncytial virus and influenza virus is characterized by the absence of pulmonary cytotoxic lymphocyte responses. J Infect Dis 195, 1126–1136 (2007).

13. Yoshida, M. et al. Local and systemic responses to SARS-CoV-2 infection in children and adults. Nature 602, 321–327 (2022).

14. Vono, M. et al. Robust innate responses to SARS-CoV-2 in children resolve faster than in adults without compromising adaptive immunity. Cell Rep 37, 109773 (2021).

15. Loske, J. et al. Pre-activated antiviral innate immunity in the upper airways controls early SARS-CoV-2 infection in children. Nat Biotechnol 40, 319–324 (2022).

16. Vareille, M., Kieninger, E., Edwards, M. R. & Regamey, N. The airway epithelium: soldier in the fight against respiratory viruses. Clin Microbiol Rev 24, 210–229 (2011).

17. Essaidi-Laziosi, M. et al. Sequential infections with rhinovirus and influenza modulate the replicative capacity of SARS-CoV-2 in the upper respiratory tract. Emerg Microbes Infect 11, 412–423 (2022).

18. Essaidi-Laziosi, M. et al. Propagation of respiratory viruses in human airway epithelia reveals persistent virus-specific signatures. J Allergy Clin Immunol 141, 2074–2084 (2018).

19. Essaidi-Laziosi, M. et al. Interferon-Dependent and Respiratory Virus-Specific Interference in Dual Infections of Airway Epithelia. Sci Rep 10, 10246 (2020).

20. Bekliz, M. et al. Immune escape of Omicron lineages BA.1, BA.2, BA.5.1, BQ.1, XBB.1.5, EG.5.1 and JN.1.1 after vaccination, infection and hybrid immunity. 2024.02.14.579654 Preprint at 10.1101/2024.02.14.579654 (2024).

21. Rameix-Welti, M.-A. et al. Visualizing the replication of respiratory syncytial virus in cells and in living mice. Nat Commun 5, 5104 (2014).

22. Baggio, S. et al. Severe Acute Respiratory Syndrome Coronavirus 2 (SARS-CoV-2) Viral Load in the Upper Respiratory Tract of Children and Adults With Early Acute Coronavirus Disease 2019 (COVID-19). Clin Infect Dis 73, 148–150 (2021).

23. Essaidi-Laziosi, M. et al. A new real-time RT-qPCR assay for the detection, subtyping and quantification of human respiratory syncytial viruses positive- and negative-sense RNAs. J Virol Methods 235, 9–14 (2016).

24. Ewels, P. A. et al. The nf-core framework for community-curated bioinformatics pipelines. Nat Biotechnol 38, 276–278 (2020).

25. Babraham Bioinformatics - FastQC A Quality Control tool for High Throughput Sequence Data. https://www.bioinformatics.babraham.ac.uk/projects/fastqc/.

26. Babraham Bioinformatics - Trim Galore! https://www.bioinformatics.babraham.ac.uk/projects/trim_galore/.

27. Patro, R., Duggal, G., Love, M. I., Irizarry, R. A. & Kingsford, C. Salmon provides fast and bias-aware quantification of transcript expression. Nat Methods 14, 417–419 (2017).

28. Ritchie, M. E. et al. limma powers differential expression analyses for RNA-sequencing and microarray studies. Nucleic Acids Res 43, e47 (2015).

29. Tarazona, S. et al. Data quality aware analysis of differential expression in RNA-seq with NOISeq R/Bioc package. Nucleic Acids Res 43, e140 (2015).

30. Robinson, M. D., McCarthy, D. J. & Smyth, G. K. edgeR: a Bioconductor package for differential expression analysis of digital gene expression data. Bioinformatics 26, 139–140 (2010).

31. Wu, T. et al. clusterProfiler 4.0: A universal enrichment tool for interpreting omics data. Innovation (Camb) 2, 100141 (2021).

32. Subramanian, A. et al. Gene set enrichment analysis: A knowledge-based approach for interpreting genome-wide expression profiles. Proceedings of the National Academy of Sciences 102, 15545–15550 (2005).

33. Wu, T. et al. clusterProfiler 4.0: A universal enrichment tool for interpreting omics data. Innovation (Camb) 2, 100141 (2021).

34. Boda, B. et al. Antiviral drug screening by assessing epithelial functions and innate immune responses in human 3D airway epithelium model. Antiviral Res 156, 72–79 (2018).

35. Hogan, B. & Tata, P. R. Cellular organization and biology of the respiratory system. Nat Cell Biol (2019) doi:10.1038/s41556-019-0357-7.

36. Ruiz García, S., et al. Novel dynamics of human mucociliary differentiation revealed by single-cell RNA sequencing of nasal epithelial cultures. Development 146, dev177428 (2019).

37. Chen, J., Kelley, W. J. & Goldstein, D. R. Role of Aging and the Immune Response to Respiratory Viral Infections: Potential Implications for COVID-19. J Immunol 205, 313–320 (2020).

38. Singanayagam, A., Joshi, P. V., Mallia, P. & Johnston, S. L. Viruses exacerbating chronic pulmonary disease: the role of immune modulation. BMC Medicine 10, 27 (2012).

39. Jonsdottir, H. R. & Dijkman, R. Coronaviruses and the human airway: a universal system for virus-host interaction studies. Virology Journal 13, 24 (2016).

40. Cao, X. et al. Invited review: human air-liquid-interface organotypic airway tissue models derived from primary tracheobronchial epithelial cells—overview and perspectives. In Vitro Cell.Dev.Biol.-Animal 57, 104–132 (2021).

41. Crystal, R. G., Randell, S. H., Engelhardt, J. F., Voynow, J. & Sunday, M. E. Airway Epithelial Cells. Proc Am Thorac Soc 5, 772–777 (2008).

42. Gras, D. et al. An *ex vivo* model of severe asthma using reconstituted human bronchial epithelium. Journal of Allergy and Clinical Immunology 129, 1259–1266.e1 (2012).

43. Balázs, A. et al. Age-Related Differences in Structure and Function of Nasal Epithelial Cultures From Healthy Children and Elderly People. Front. Immunol. 13, (2022).

44. Woodall, M. N. J. et al. Age-specific nasal epithelial responses to SARS-CoV-2 infection. Nat Microbiol 9, 1293–1311 (2024).

45. Smith, C. M. et al. Ciliary dyskinesia is an early feature of respiratory syncytial virus infection. European Respiratory Journal 43, 485–496 (2014).

46. Essaidi-Laziosi, M. et al. Altered cell function and increased replication of rhinoviruses and EV-D68 in airway epithelia of asthma patients. Front Microbiol 14, 1106945 (2023).

47. Maughan, E. F. et al. Cell-intrinsic differences between human airway epithelial cells from children and adults. iScience 25, 105409 (2022).

48. Watkins, T. A. et al. High burden of viruses and bacterial pathobionts drives heightened nasal innate immunity in children. J Exp Med 221, e20230911 (2024).

49. Magalhães, V. G. et al. Immune-epithelial cell cross-talk enhances antiviral responsiveness to SARS-CoV-2 in children. EMBO Rep 24, e57912 (2023).

50. Zimmermann, P. & Curtis, N. Why Does the Severity of COVID-19 Differ With Age?: Understanding the Mechanisms Underlying the Age Gradient in Outcome Following SARS-CoV-2 Infection. Pediatr Infect Dis J 41, e36–e45 (2022).

51. Pierce, C. A. et al. Natural mucosal barriers and COVID-19 in children. JCI Insight 6, (2021).

52. Mick, E. et al. Upper airway gene expression shows a more robust adaptive immune response to SARS-CoV-2 in children. Nat Commun 13, 3937 (2022).

53. Koch, C. M. et al. Age-related Differences in the Nasal Mucosal Immune Response to SARS-CoV-2. Am J Respir Cell Mol Biol 66, 206–222 (2022).

54. Chang, J. J.-Y. et al. Uncovering strain- and age-dependent innate immune responses to SARS-CoV-2 infection in air-liquid-interface cultured nasal epithelia. iScience 27, 110009 (2024).

55. Stölting, H. et al. Distinct airway epithelial immune responses after infection with SARS-CoV-2 compared to H1N1. Mucosal Immunol 15, 952–963 (2022).

56. Garofalo, R. P. et al. A comparison of epidemiologic and immunologic features of bronchiolitis caused by influenza virus and respiratory syncytial virus. J Med Virol 75, 282–289 (2005).

57. Usemann, J., Alves, M. P., Ritz, N., Latzin, P. & Müller, L. Age-dependent response of the human nasal epithelium to rhinovirus infection. European Respiratory Journal 56, (2020).

58. O Loughlin, D. W., Coughlan, S., De Gascun, C. F., McNally, P. & Cox, D. W. The role of rhinovirus infections in young children with cystic fibrosis. J Clin Virol 129, 104478 (2020).

59. Jackson, D. J. & Gern, J. E. Rhinovirus Infections and Their Roles in Asthma: Etiology and Exacerbations. J Allergy Clin Immunol Pract 10, 673–681 (2022).

60. Gern, J. E. & Busse, W. W. Association of Rhinovirus Infections with Asthma. Clin Microbiol Rev 12, 9–18 (1999).

61. Principi, N. et al. Prospective evaluation of rhinovirus infection in healthy young children. J Clin Virol 66, 83–89 (2015).

62. Chen, W.-J. et al. Epidemiologic, clinical, and virologic characteristics of human rhinovirus infection among otherwise healthy children and adults. J Clin Virol 64, 74–82 (2015).

63. Bellon, M. et al. Severe Acute Respiratory Syndrome Coronavirus 2 (SARS-CoV-2) Viral Load Kinetics in Symptomatic Children, Adolescents, and Adults. Clin Infect Dis 73, e1384–e1386 (2021).

64. Reina, J., Morales, C., Busquets, M. & Norte, C. Usefulness of Ct value in acute respiratory infections caused by respiratory syncytial virus A and B and influenza virus A (H1N1)pdm09, A (H3N2) and B. Enferm Infecc Microbiol Clin (Engl Ed) 36, 332–335 (2018).

65. Kaler, J., Hussain, A., Patel, K., Hernandez, T. & Ray, S. Respiratory Syncytial Virus: A Comprehensive Review of Transmission, Pathophysiology, and Manifestation. Cureus 15, e36342.

66. Peltola, V., Waris, M., Kainulainen, L., Kero, J. & Ruuskanen, O. Virus shedding after human rhinovirus infection in children, adults and patients with hypogammaglobulinaemia. Clinical Microbiology and Infection 19, E322–E327 (2013).

67. Ng, S. et al. The Timeline of Influenza Virus Shedding in Children and Adults in a Household Transmission Study of Influenza in Managua, Nicaragua. Pediatr Infect Dis J 35, 583–586 (2016).

68. Sato, M., Hosoya, M., Kato, K. & Suzuki, H. VIRAL SHEDDING IN CHILDREN WITH INFLUENZA VIRUS INFECTIONS TREATED WITH NEURAMINIDASE INHIBITORS. The Pediatric Infectious Disease Journal 24, 931 (2005).

69. Fielding, J. E., Kelly, H. A., Mercer, G. N. & Glass, K. Systematic review of influenza A(H1N1)pdm09 virus shedding: duration is affected by severity, but not age. Influenza Other Respir Viruses 8, 142–150 (2014).

70. Han, M. S. et al. Clinical Characteristics and Viral RNA Detection in Children With Coronavirus Disease 2019 in the Republic of Korea. JAMA Pediatr 175, 73–80 (2021).

