## supplementary tables and figures for "Age- and Virus-Specific Signatures of *In Vitro* Reconstituted Human Airway Epithelia in the Presence and Absence of Respiratory Viral Infections"

**Table S1: Donor information**

| Anonymized participants | age group | age | sex | family | family link |
|---|---|---|---|---|---|
| CH1 | child | 9 | male | 1 | son |
| CH2 | child | 5 | female | 1 | daughter |
| CH3 | child | 10 | female | 2 | son |
| CH4 | child | 8 | female | 2 | daughter |
| CH6 | child | 6 | male | 3 | son |
| CH7 | child | 9 | male | 4 | son |
| CH9 | child | 9 | female | 5 | daughter |
| CH13 | child | 6 | female | 7 | daughter |
| CH15 | child | 6 | male | 8 | son |
| CH17 | child | 6 | female | 9 | daughter |
| A2 | adult | 50 | female | 2 | mother |
| A3 | adult | 27 | female | 2 | sister |
| A4 | adult | 39 | female | 1 | mother |
| A7 | adult | 29 | female | 10 | NA |
| A11 | adult | 34 | female | 5 | mother |
| A14 | adult | 38 | female | 7 | mother |
| A15 | adult | 41 | female | 7 | father |
| A17 | adult | 43 | female | 4 | mother |
| A20 | adult | 37 | female | 9 | mother |

**Table S2: Virus stock information**

| Virus | | Sequences | Origin | Titration | MOI | Origin/reference |
|---|---|---|---|---|---|---|
| SARS-CoV-2 | EG.5.1 | hCoV-19/Switzerland/GE-HUG-42157557/2023 (in GISAID) | Clincal sample isolated in Vero E6 | In Vero E6-TMPRSS2 cells | 0.01 | [1] |
| IAV | H1N1 pan2009 | Not available | Clinical sample isolated in MucilAir™ | In MDCK cells | 0.002 | [2] |
| RV | RV-A16 | Not available | Clinical sample isolated in MucilAir™ | In MucilAir™ | 0.002-0.006 | This study |
| RV | RV-A16 | Not available | Lab-adapted virus (ATCC) | In MucilAir™ | 0.0035 | Purchased from ATCC |
| RSV | RSV-A | Not available | Clinical sample isolated in MucilAir™ | In MucilAir™ | 0.018 | This study |

| | | | | | | |
|---|---|---|---|---|---|---|
| RSV | RSV-A mCherry | Not available | Recombinant strain | In MucilAir™ | 0.01 | [3] |

#### Table S3: References of the selected genes of interest for the heatmaps

| Gene related to | List of selected genes | References |
|---|---|---|
| Basal cells | KRT5, ITGB6, DLK2, KRT14, KRT15, TP63, DAPL1, Notch1, Notch3, COL3A1, MKI67, NUSAP1, LY6D, KRT6A, KRT13, BCAM, VIM, EPCAM, CDH1, FN1, COL1A1, CDH2, TNC, VCAN, PCP4, CUX2, SPINK1, PRSS2, CPA6, CTSE, MMP7, MDK, GDF15, PTGS2, SLCO2A1, EPHB2, ITGB8, ITGAV, ITGB6, TGFB1, KCNN4, KCNQ5, KCNS3, CDKN1A, CDKN2A, CDKN2B, CCND1, CCND2, MDM2, HMGA2, PTCHD4, OCIAD2 | [4,5] |
| Goblet cells | MUC1, MUC12, MUC13, MUC15, MUC16, MUC2, MUC20, MUC21, MUC22, MUC3A, MUC4, MUC5AC, MUC5B, MUCL1, SCGB1A1, TFF3, TFF1, BPIFA1, PLAU, SCGB3A1, NOS2, CAPN13, RARRES1, SERPINB3, AQP5, CP, PI3, SPRR3, KRT78, FP671120.4, FP236383.2, PIGR, VMO1, XBP1, CYP2B6, CYP2A13 | [4–10] |
| Ciliated cells | FOXJ1, 2DEFB1, SPTBN5, EHD1, USH1G, RP1L1, CCDC68, MYRIP, GSTM3, CLTB, GPR157, CCDC120, CIB2, CYLD, RP2, PSEN1, PTPRK, RAB8A, AMBRA1, SCLT1, CCSAP, KLC3, CDKL5, EYS, MARK4, APP, AKT1, CSNK1A1, SQSTM1, LYAR, GPR37L1, AK2, GNB1, PROM2, PRKACA, PKD1L1, EPS15, TUBG1, ATG7, TBCC, PTPN23, GLI3, RAB27A, MAP4, MERTK, ARFGEF2, HYAL3, DNAH17, SLC26A6, DYNLL2, SAXO1, PRKAR1B, DCTN1, OCRL, KIF5B, SHANK3, OFD1, ATG5, ATP2B4, KIF7, RABEP2, FOPNL, SMO, MAP1LC3B, WDR11, PIK3C3, ATG16L1, CCDC88A, SCNN1A, WRAP73, TMEM216, GNA11, DHRS3, KIF3C, DLG5, DAAM1, EVC2, ATG14, SSNA1, PIK3R4, POC1A, TOPORS, MYO5A, CENPJ, CASK, SUFU, CCDC66, PRKAR1A, TBC1D30, CENPF, TAPT1, CEP164, DYNLRB1, TCP11, SSX2IP, SNAP29, DDX6, SLC9B2, RTTN, BBS7, ARHGAP35, EVC, ERC1, NEK8, ICK, DYNLL1, BBIP1, CNGB1, GRK4, EZR, PKD1, TMEM237, TCTN3, KIF2A, TTC30B, FAM161A, GABARAP, CDK10, NEK4, PCDHB13, TMEM17, CEP170, PDE6B, GLI1, SLC9A3R1, B9D2, CFAP36, RAB28, DCDC2, CATSPERG, IFT43, MAK, MXRA8, TCTEX1D2, PTCH1, C8orf37, EHD3, IFT27, INPP5E, KIAA0586, C2CD3, ODF2L, POC1B, TBC1D7, CFAP46, GNAQ, AKAP3, NAPEPLD, TTLL6, IFT46, CEP41, TULP3, USH2A, TRAF3IP1, DYNC2LI1, CFAP20, ODF2, DNAH5, CFAP126, CEP131, RSPH1, MOK, PKD2, WDR34, GAS8, RSPH9, MAP1B, KIF3B, PRKAR2A, RABL2B, FBF1, ULK3, ROM1, TCTN1, CEP290, AHI1, SPEF1, IFT20, DNALI1, IQCE, DZIP1, MAPK15, PCDHB15, AK8, DNAH2, KIF19, PRCD, TTBK2, PCM1, INTU, CCDC65, RGS9BP, WDR35, WDR60, BBS1, ARL3, CNGA4, TTC8, TTC26, IFT57, WDPCP, IQCB1, CCDC40, IFT22, KIF3A, CFAP61, LCA5, CNTRL, CCDC151, ARL6, MKS1, C5orf30, CEP19, RSPH4A, GPR161, BBS5, RPGR, BBS2, IFT172, PRKACB, NPHP4, UNC119B, PIFO, ENKUR, DNAH3, IFT74, IFT122, CEP89, DYNLRB2, CEP78, PRKAR2B, KIFAP3, TCTEX1D4, IFT140, SPATA6, BBS4, SPA17, CEP128, IFT52, HSPB11, DNAJB13, B9D1, DNAL1, SPATA7, TEKT4, CFAP54, NPHP1, CLUAP1, ARL13B, USH1C, CEP126, RP1, TMEM231, TTC21B, SPAG6, DNAH9, CCDC114, MLF1, CC2D2A, AGBL4, DNAAF1, CETN2, SHANK2, PACRG, DNAH1, ABCA4, IFT88, WDR19, WDR66, DRD2, CFAP221, CFAP74, DRD1, IFT81, NIN, DNAI2, ARMC9, PROM1, CCDC39, AGBL2, AKAP14, IFT80, CEP162, CCDC103, EFCAB7, RPGRIP1L, BBS9, JADE1, TMEM67, CDC14A, MAATS1, TMEM107, CEP83, MNS1, SPAG16, DNAI1, DNAH6, HYDIN, FANK1, DNAH7, KIF17, PEX6, CERKL, ARMC4, TACR1, TCTN2, DZIP1L, EFHC1, CROCC, TTLL3, CFAP69, CDHR1, CFAP43, CATSPERD, CDKL1, GLI2, C11orf97, DYNC2H1, CACNA1F, PDE6A, GUCA1B, DRC1, CABYR, DNAH11, PRPH2, OMG, FOXN4, CDC20B, FOXI1, ASCL3, SAA4, SAA2, SAA1, CCDC113, CCDC153, MYCL, CCNO | [4–7]<br>Genego ontologies |
| Interferon response | IRF3, DHX58 (LGP2), IFNB1, IFNL1, IFNL2, IFNL3, IFNG, IFNA1, IFIT1, CXCL11, C1orf29, ISG15, IFI27, Viperin, CXCL10, MX1, MX2, G1P3, OAS2, IRF7, OAS1, IFITM1, IFIT4, GBP1, WARS, IFIT2, IFI35, STAT1, IFI44, ISG20, OASL, PKR, ISGF3G, IFITM2, IFITM3, IFRG28, USP18, SP110, LAMP3, BST2, FLJ20637, LOC51191, HSXIAPAF1, LAP3, FLJ20073, FLJ22693, CIC, FLJ20035, APOBEC3A, HRASLS2, TOR1B, ECGF1, TRIM14, APOBEC3, | [4,5,11,12] |

| | | |
|---|---|---|
| | BST2 (tetherin), C6orf150 (MB21D1), CD74, DDIT4, DDX58 (RIG-I), DDX60, EIF2AK2 (PKR), GBP1, GBP2, HPSE, IFI44L, IFI6/G1P3, IFIH1 (MDA5), IFIT1/2/3/5, IFITM1/2/3, IRF1, MAP3K14 (NIK), MOV10, MS4A4A, MX1 (MxA), MX2 (MxB), NAMPT (PBEF1), NT5C3, OAS1/2/3, P2RY6, PHF15, PML (TRIM19), RSAD2 (viperin), RTP4, SLC15A3, SLC25A28, SSBP3, TREX1 (AGS1), TRIM5, TRIM25, SUN2 (UNC84B), ZC3HAV1 (ZAP), RSAD2, IRF9, IFNAR2, STAT2, TYK2, JAK1, STAT3, IFNAR1, ADAR, AXL, BST2, EIF2AK2, GAS6, GATA3, IFIT2, IFIT3, IFITM1, IFITM2, IFITM3, IFNAR1, IFNAR2, KLHL20, LAMP3, MX2, PYHIN1, RO60, TPR, OAS3, OASL, OTOP1, PARP14, PARP9, PDE12, PIAS1, PML, PPARG, PRKCD, PTAFR, PTPN2, RAB20, RAB43, RAB7B, RPL13A, RPS6KB1, SHFL, SIRPA, SLC11A1, SLC26A6, SLC30A8, SNCA, SOCS1, SOCS3, SP100, STAR, STAT1, STX4, STX8, STXBP1, STXBP3, STXBP4, SUMO1, SYNCRIP, TDGF1, TLR2, TLR3, TLR4, TP53, TRIM21, TRIM22, TRIM25, TRIM26, TRIM31, TRIM34, TRIM38, TRIM5, TRIM62, TRIM68, TRIM8, TXK, UBD, VAMP3, VCAM1, VIM, VPS26B, WAS, WNT5A, XCL1, XCL2, ZYX, ZBP1, MSR1, C9orf91, IFIT5, LY6E | |
| Inflammation | IL1a, IL1b, IL1raP, IL2Rb, IL5Ra, IL6, IL10Rb, IL10RB_AS1, IL11, IL11Ra, IL12Rb1, IL12Rb2, IL13Ra1, IL16, IL32, IL3Ra, LIF, LIFR, LIFR_AS1, LTB4R, LTB4R2, LTBP1, LTBP2, LTBP3, LTBP4, OSMR, TGFb1, TGFb2, TGFb3, TNFRSF1A, TNFRSF1B, TSLP, IL17B, IL17C, IL17D, IL33, IL18, IL27, CXCL10 | [13–16] |

### Supplementary figures:

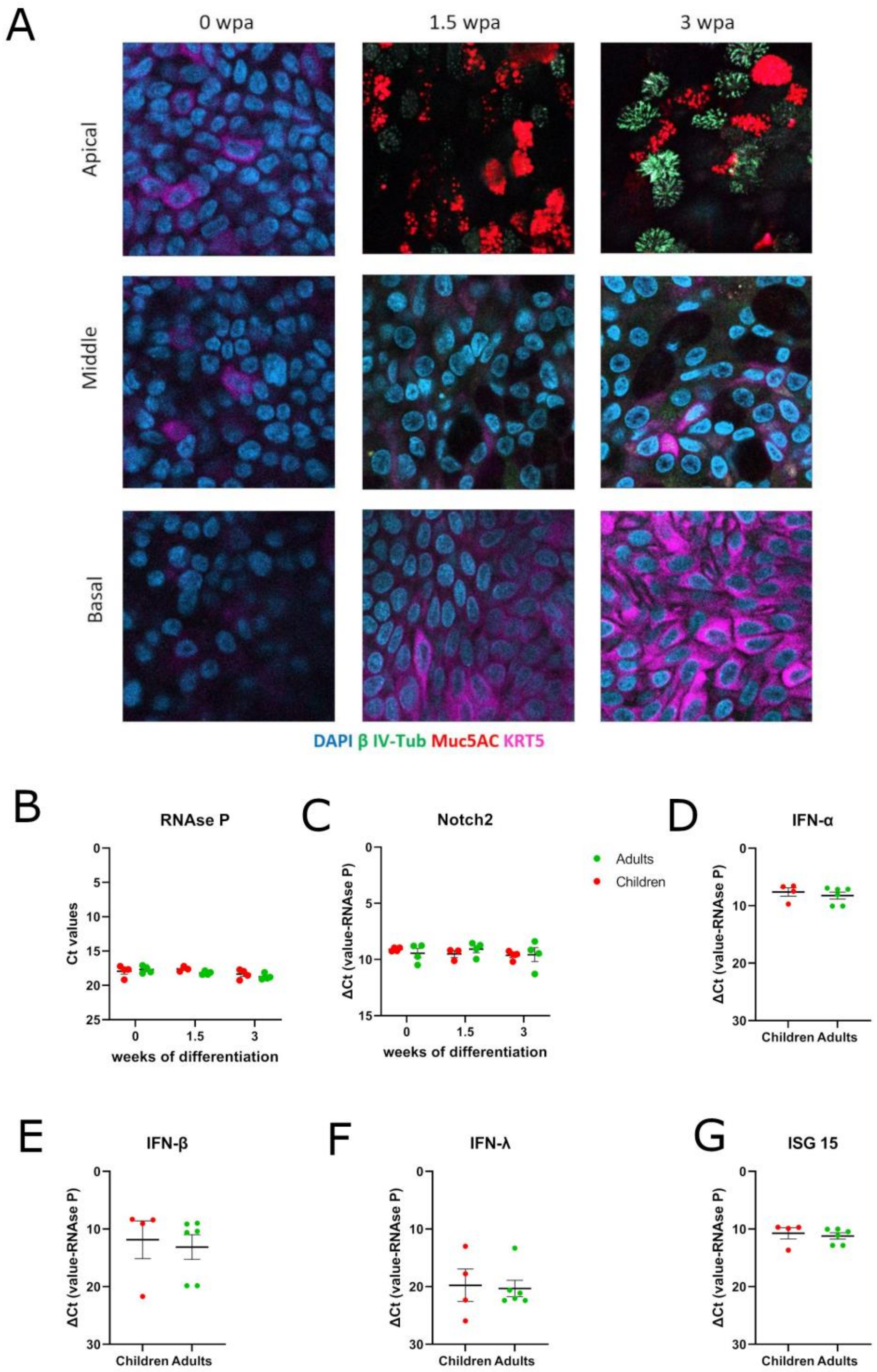


**Supplementary Figure S1: Additional characterization of the in-house *in vitro* differentiated HAE. A:** Immunofluorescence of a child tissue, as an example, at different stages of differentiation and at 3 different section levels (apical, middle, and basal) of the epithelial tissue. Blue: DAPI (nuclei), green: β IV-tubulin (ciliated cells), red: mucin 5AC (goblet cells), pink: cytokeratin 5 (basal cells). Images were acquired with Zeiss LSM700 Meta Confocal Microscope. **B, C:** From the same experiments as in the figure 1, expression of RNAse P (B), Notch 2 (C), in tissues from children (N=4) and adults (N=4) at different stages of differentiation measured by RT-PCR. **D, E, F, G:** Expression of IFN-α (D), -β (E), -λ (F) and ISG15 (G) in non-infected HAE from children (N=4) and adults (N=6) measured by RT-PCR at 3wpa (after full differentiation).

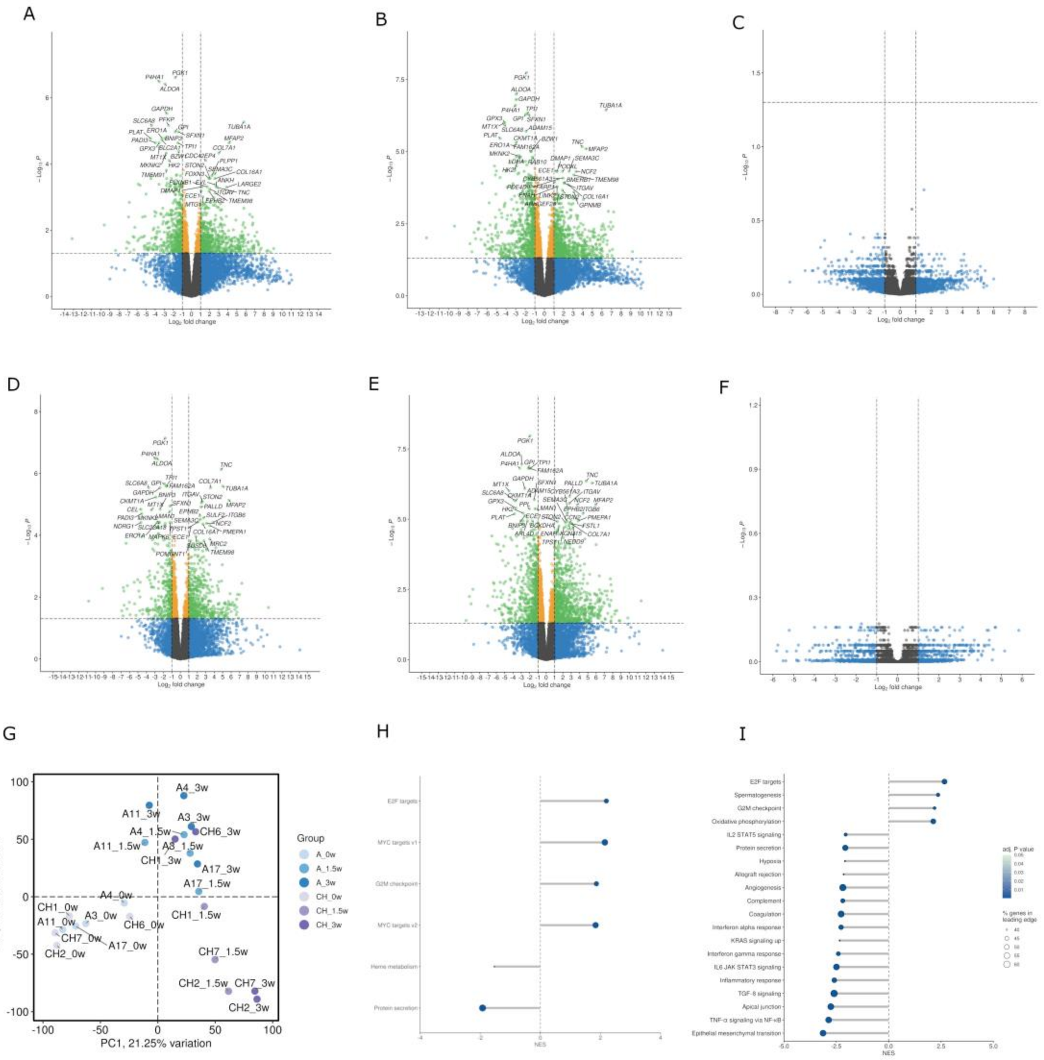


**Supplementary figure S2: Differential gene expression in HAE from children versus adults during *in vitro* differentiation. A, B, and C:** Volcano plots of differential gene expression between 0 and 1.5 wpa (A), 0 and 3 wpa (B), 1.5 and 3 wpa (C) in tissues from children. **D, E, and F:** Volcano plots of differential gene expression between 0 and 1.5 wpa (D), 0 and 3 wpa (E), 1.5 and 3 wpa (F) in adults' tissues. $Log_2$ fold change ($Log_2$ FC) threshold=1. Adjusted P value threshold=0.05. Grey dots: not significant (NS); green dots: $Log_2$ FC <-1 or >1 and P value >0.05; blue dots: only $Log_2$ FC <-1 or >1; orange dots: only P value >0.05. **G:** Principal component analysis (PCA) plot of all differentiation samples N adults=4, N children=4. **H and I**: Hallmark gene set enrichment analysis (GSEA) of child tissues compared to adult tissues at 0 wpa (H) and 1.5 wpa (I). Positive normalized enrichment score (NES): pathways upregulated in children, negative NES: pathways upregulated in adults. Size of the dots: percentage of genes in the leading edge (from 10 to 40%). Color of the dots: adjusted P value (from 0.01 (darkest blue) to 0.05 (lightest blue)).

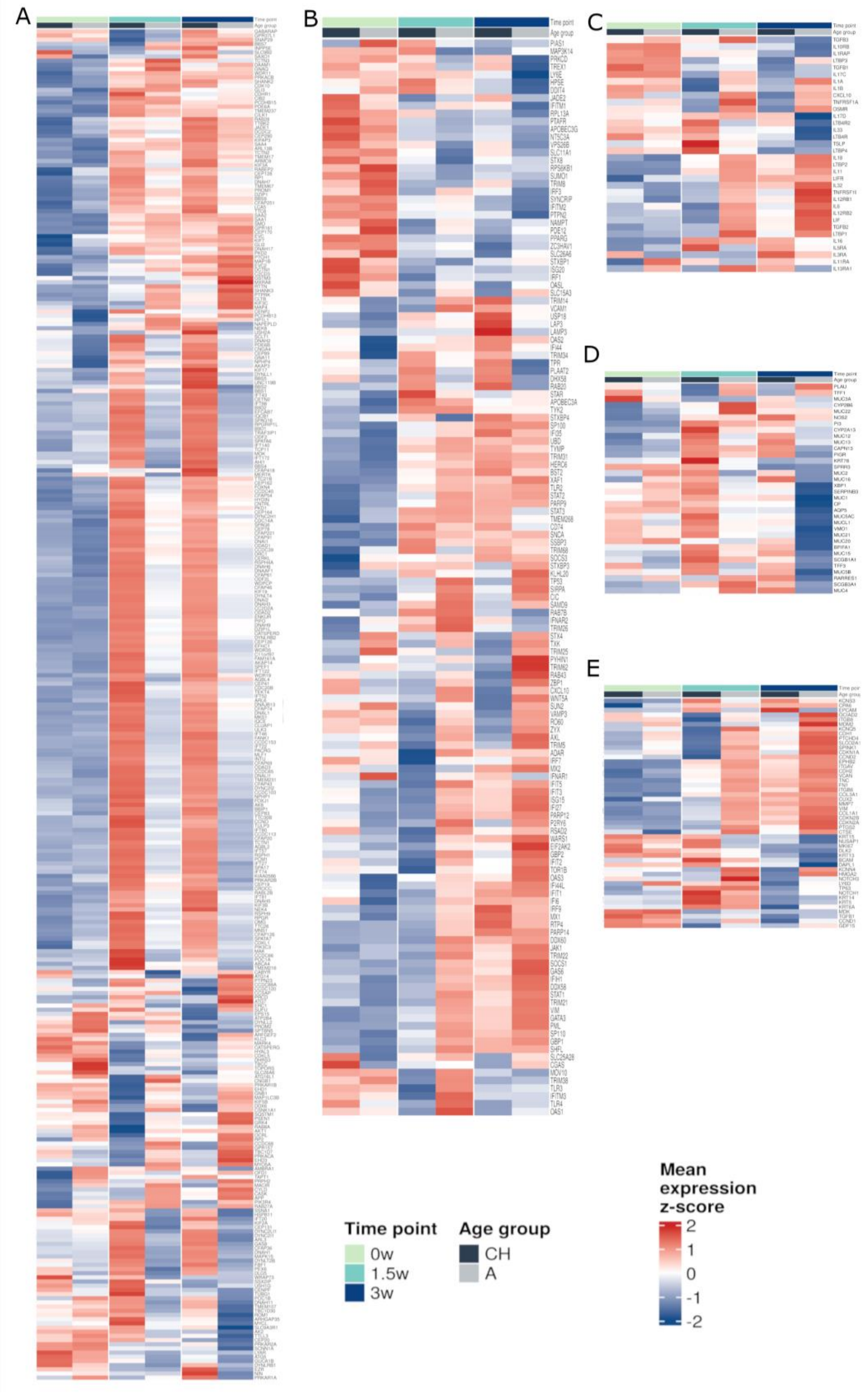


**Supplementary figure S3: HAE gene expression during the differentiation.** Heatmaps of selected genes related to ciliated cells (A), interferon signalling pathways (B), inflammation (C), goblet cells (D), and basal cells (E) in child (N=4) and adult (N=4) tissues at 0, 1.5, and 3wpa.

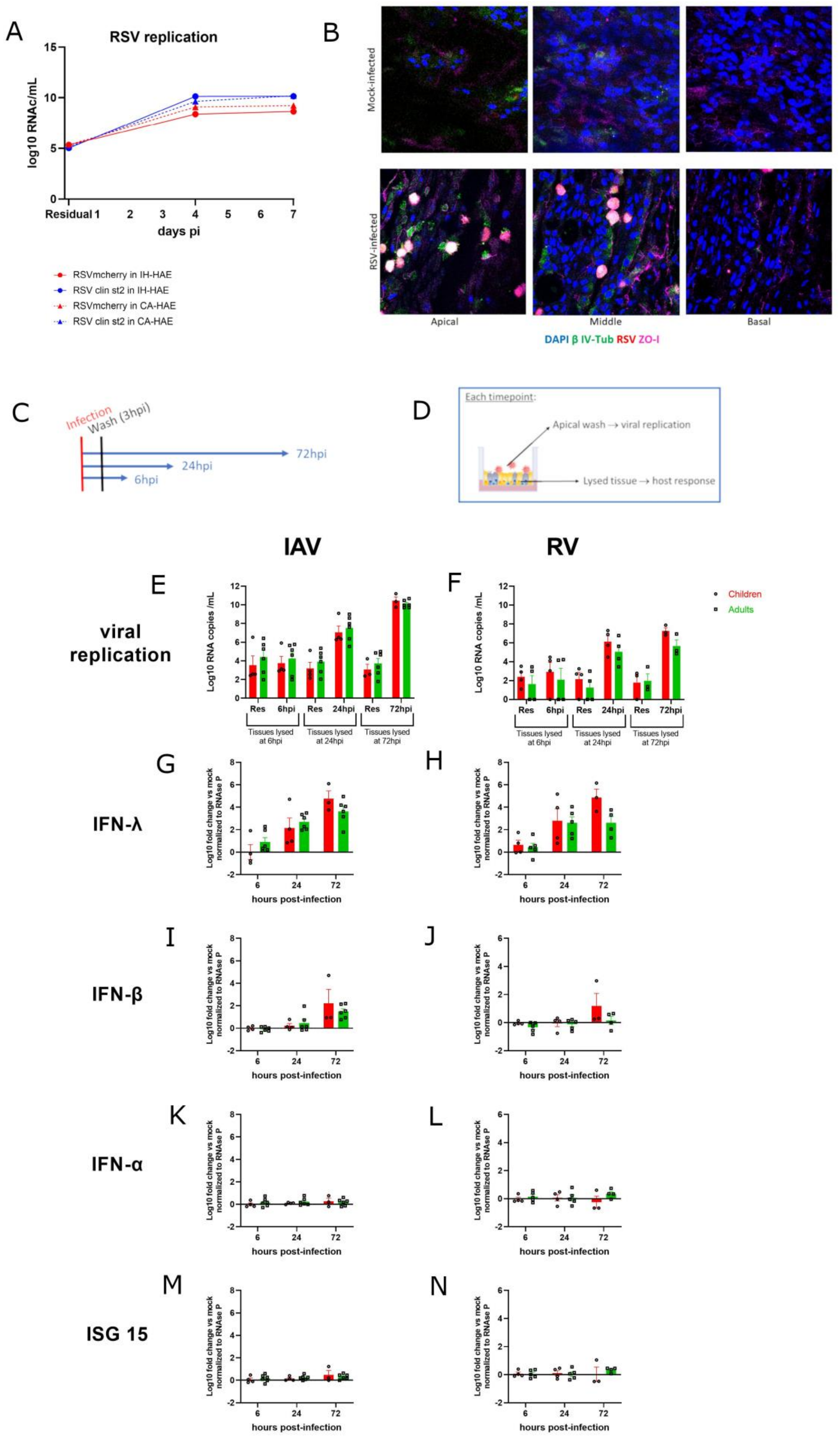

A
RSV replication
log10 RNAc/mL
days pi
Residual 1
RSVmcherry in IH-HAE
RSV clin st2 in IH-HAE
RSVmcherry in CA-HAE
RSV clin st2 in CA-HAE
B
Mock-infected
RSV-infected
Apical
Middle
Basal
DAPI β IV-Tub RSV ZO-I
C
Infection
Wash (3hpi)
72hpi
24hpi
6hpi
D
Each timepoint:
Apical wash → viral replication
Lysed tissue → host response
IAV
RV
E
F
viral replication
Log10 RNA copies /mL
Res
6hpi
24hpi
72hpi
Tissues lysed at 6hpi
Tissues lysed at 24hpi
Tissues lysed at 72hpi
Children
Adults
G
H
IFN-λ
I
J
IFN-β
K
L
IFN-α
M
N
ISG 15
Log10 fold change vs mock normalized to RNAse P
hours post-infection

**Supplementary figure S4: Validation of the in-house model as tool for respiratory viral infection A and B.** RSV infection in an in-house adult tissue and in commercially available nasal tissue. **A:** Viral replication of RSV A mCherry and clinical isolate of RSV A (RSVclin) in in-house adult tissue (IH-HAE) and in commercially available nasal tissue (CA-HAE, Epithelix Mucilair) measured by RT-PCR. The infection was performed at 37°C with a MOI of 0.1 for RSV A mCherry and a MOI of 0.018 for the clinical RSVA. **B:** Immunofluorescence of mock-infected and RSV-A mCherry infected in-house adult tissue at 3 different levels: apical, middle and basal part of the tissue. Blue: DAPI (nuclei), green: β-tubulin (ciliated cells), red: mucin 5AC (muc5AC, goblet cells) in mock, red: RSV A mCherry in RSV-infected tissues. Images were acquired with Zeiss LSM700 Meta Confocal Microscope. **C to N**. Pilot infection assay to study viral replication and host response in the context of IAV and RV infections. **C and D:** Design of the infection experiment: the child and adult tissues were infected at 33°C with diluted clinical stocks of either IAV (H1N1) or RV (RV-A16). After 3 hours of infection (3hpi), the tissues were washed and the residual apical wash was collected. At each timepoint, 6, 24 and 72hpi, apical washes were collected, to quantify the viral replication, and tissues were lysed, to measure the host response. **E and F**: IAV (E) and RV (F) replication in the apical washes of the lysed tissues compared to a residual apical wash collected on the same tissue. Measured by RT-PCR. MOI=0.002 for IAV and 0.002-0.006 for RV. N children=4, N adults=6. Mean and standard deviation are represented. No statistically significant difference between children and adults was observed. **G to N**: IFN-λ (G and H), -β (I and J), α (K and L) and ISG15 (M and N) induction, in the context of IAV or RV infection, measured by RT-PCR in total intracellular RNA from tissue lysates, expressed in fold change compared to the mock, normalized with the expression level of RNAseP. Mean and standard deviation are represented. No statistically significant difference between children and adults was measured.

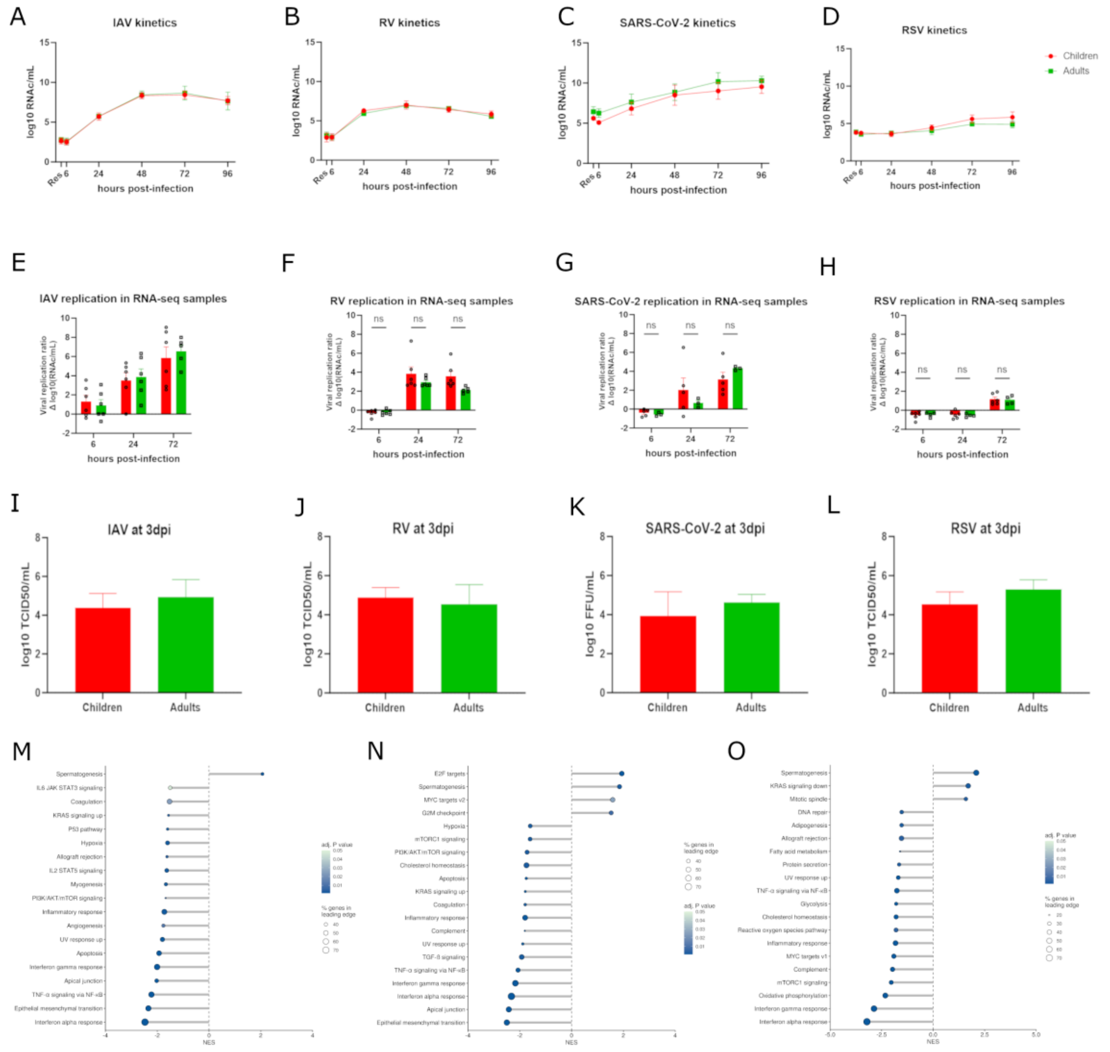


**Supplementary figure S5: Viral replication kinetics of Influenza A, Rhinovirus A16, SARS-CoV-2 Omicron EG5.1 and RSV A from the assays used for the transcriptomic study. A:** Influenza A H1N1 virus (IAV) replication kinetics in child and adult tissues measured by RT-qPCR. The infection was performed at 33°C, MOI 0.02. N children=7, N adults=5. **B:** Rhinovirus A16 (RV) replication kinetics in child and adult tissues measured by RT-qPCR. The infection was performed at 33°C, MOI 0.0035. N children=5, N adults=4. **C:** SARS-CoV-2 Omicron EG.5.1 replication kinetics in child and adult tissues measured by RT-qPCR. The infection was performed at 33°C, MOI 0.01. N children=4, N adults=3. Means and standard deviations are represented. **D:** RSV-A mCherry replication kinetics. The infection was performed at 33°C, MOI 0.01. N children=3, N adults=3. Means and standard deviations are represented **E, F, G, H:** Viral replication of IAV (N children=6, N adults=6) (E), RV (N children=6, N adults=6) (F), SARS-CoV-2 (N children=5, N adults=3) (G) and RSV (N children=6, N adults=4) (H). Viral replication was measured in the inserts used for the transcriptomic analysis and is expressed as a ratio to the value measured in the same insert at 3hpi (residual) in Δ log10(RNAc/mL). Infections performed in the same conditions as in A, B, C, D. **I.J.K.L**: viral titers at 3dpi of IAV (N children=6, N adults=6) (I), RV (N children=6, N adults=6) (J), SARS-CoV-2 (N children=5, N adults=3) (K) and RSV (N children=6, N adults=4) (L) determined by assessing the Tissue Culture Infectious Dose 50 (TCID50) for IAV, RSV and RV and by calculating the Focus forming Units (FFU) for SARS-CoV-2, as previously described[2,17,18]. **M, N, O:** Hallmark gene set enrichment analysis (GSEA) of mock-infected child tissue samples (N=9) compared to adults (N=6) collected at 6hpi (M), 24hpi (N), and 72hpi (O). Positive normalized enrichment score (NES): pathways upregulated in children, negative NES: pathways upregulated in adults. Size of the dots: percentage of genes in the leading edge. Colour of the dots: adjusted P value.

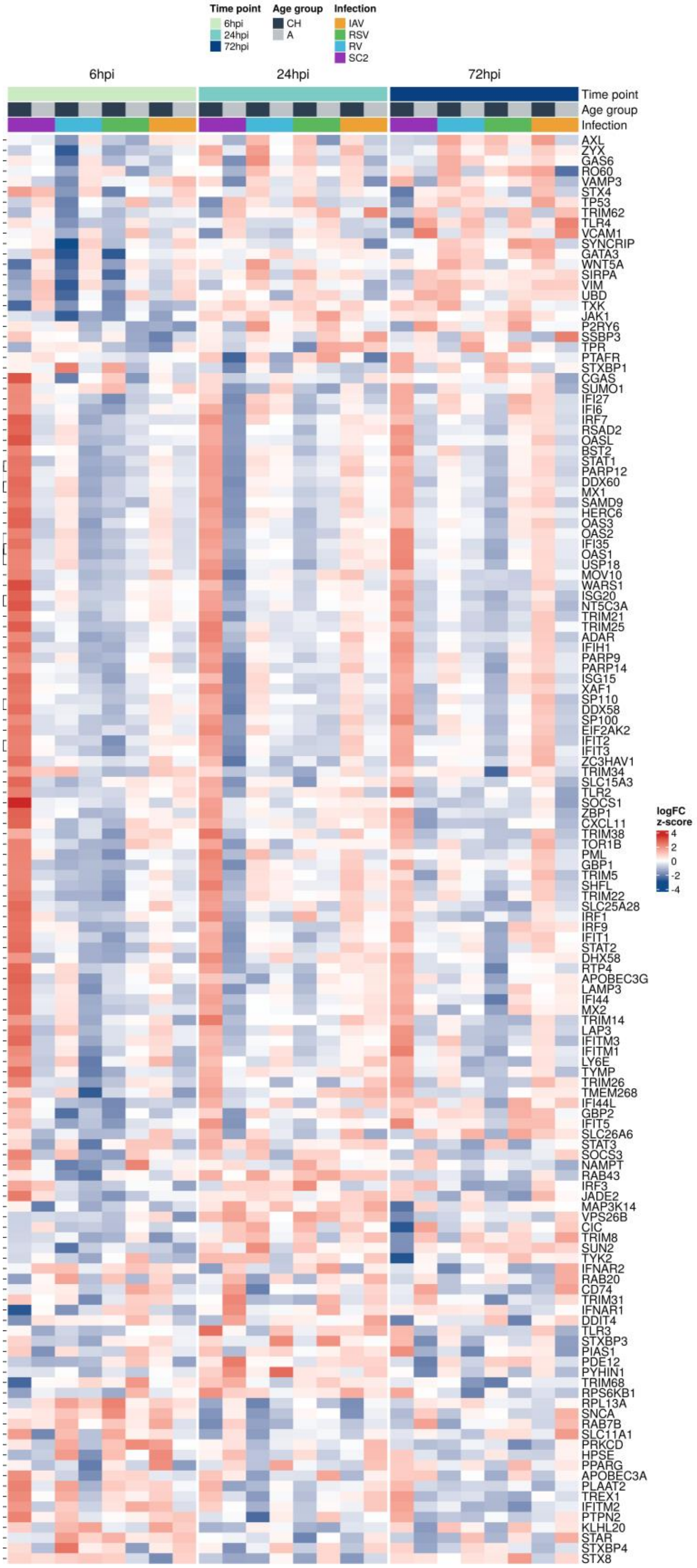


**Supplementary figure S6: Detailed heatmap of selected genes involved in interferon signalling pathway during infection by SARS-CoV-2, RV, RSV, and IAV in children and adults** (N: similar as in figure 5 for the different viruses) at 6hpi, 24hpi, 72hpi, normalized with the gene expression in the corresponding mock tissue.

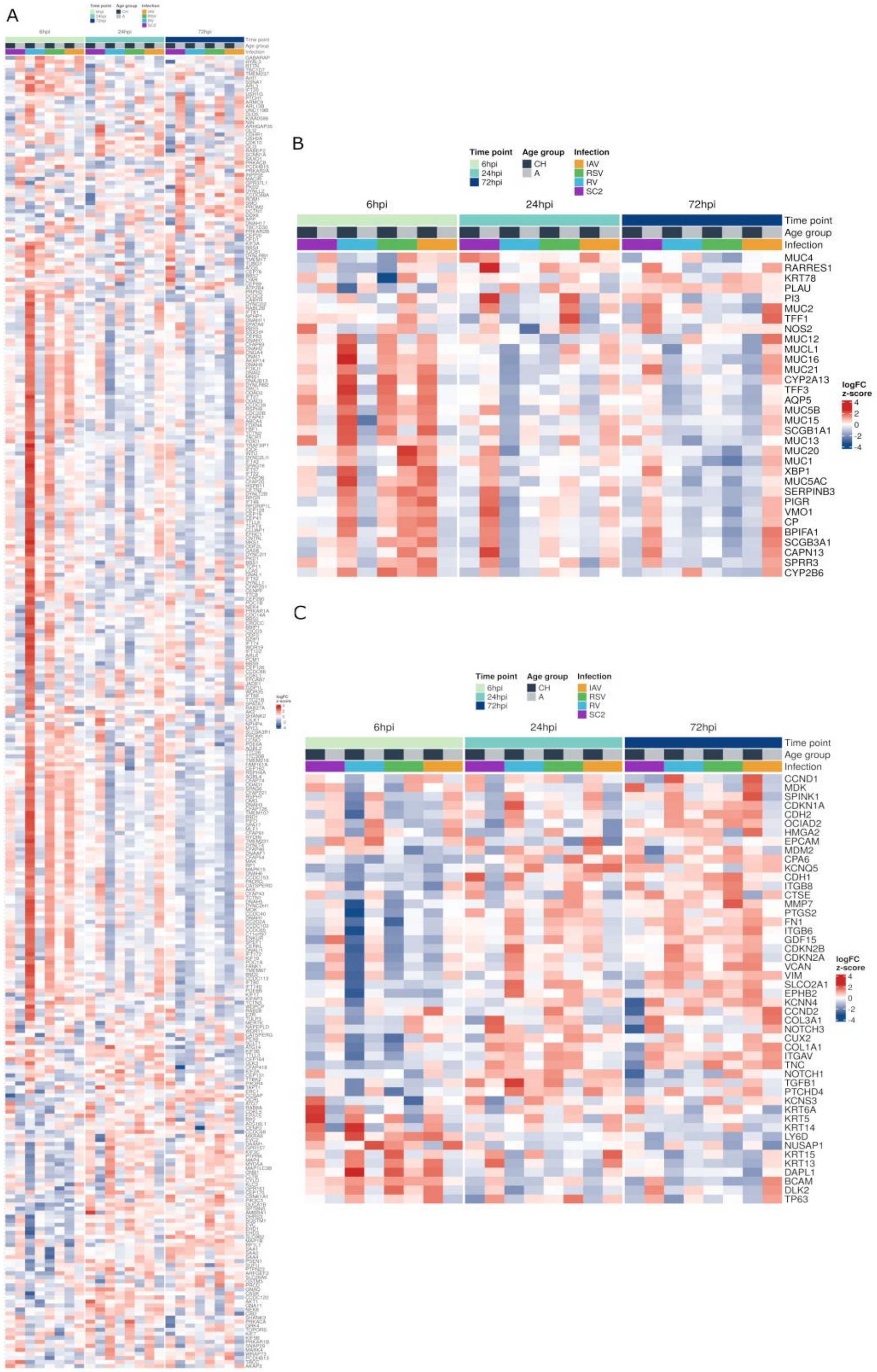


**Supplementary figure S7: Differential expression of genes to epithelial cells in HAE from children versus adults infected by SARS-CoV-2, RV, RSV and IAV at 6hpi, 24hpi, and 72hpi**: Heatmaps of selected genes related to ciliated cells (A), goblet cells (B), and basal cells (C) in tissues infected by SARS-CoV-2, RV, RSV, and IAV, normalized with the gene expression in the corresponding mock tissue, in children and adults (N: similar as in figure 5 for the different viruses) at 6hpi, 24hpi, 72hpi.

### Supplementary references:


(1) Bekliz, M.; Essaidi-Laziosi, M.; Adea, K.; Hosszu-Fellous, K.; Alvarez, C.; Bellon, M.; Sattonnet-Roche, P.; Puhach, O.; Dbeissi, D.; Zaballa, M. E.; Stringhini, S.; Guessous, I.; Vetter, P.; Eberhardt, C. S.; Kaiser, L.; Eckerle, I. Immune Escape of Omicron Lineages BA.1, BA.2, BA.5.1, BQ.1, XBB.1.5, EG.5.1 and JN.1.1 after Vaccination, Infection and Hybrid Immunity. bioRxiv February 15, 2024, p 2024.02.14.579654. https://doi.org/10.1101/2024.02.14.579654.

(2) Essaidi-Laziosi, M.; Alvarez, C.; Puhach, O.; Sattonnet-Roche, P.; Torriani, G.; Tapparel, C.; Kaiser, L.; Eckerle, I. Sequential Infections with Rhinovirus and Influenza Modulate the Replicative Capacity of SARS-CoV-2 in the Upper Respiratory Tract. *Emerg Microbes Infect* **2022**, *11* (1), 412–423. https://doi.org/10.1080/22221751.2021.2021806.

(3) Rameix-Welti, M.-A.; Le Goffic, R.; Hervé, P.-L.; Sourimant, J.; Rémot, A.; Riffault, S.; Yu, Q.; Galloux, M.; Gault, E.; Eléouët, J.-F. Visualizing the Replication of Respiratory Syncytial Virus in Cells and in Living Mice. *Nat Commun* **2014**, *5*, 5104. https://doi.org/10.1038/ncomms6104.

(4) Woodall, M. N. J.; Cujba, A.-M.; Worlock, K. B.; Case, K.-M.; Masonou, T.; Yoshida, M.; Polanski, K.; Huang, N.; Lindeboom, R. G. H.; Mamanova, L.; Bolt, L.; Richardson, L.; Cakir, B.; Ellis, S.; Palor, M.; Burgoyne, T.; Pinto, A.; Moulding, D.; McHugh, T. D.; Saleh, A.; Kilich, E.; Mehta, P.; O'Callaghan, C.; Zhou, J.; Barclay, W.; De Coppi, P.; Butler, C. R.; Cortina-Borja, M.; Vinette, H.; Roy, S.; Breuer, J.; Chambers, R. C.; Heywood, W. E.; Mills, K.; Hynds, R. E.; Teichmann, S. A.; Meyer, K. B.; Nikolić, M. Z.; Smith, C. M. Age-Specific Nasal Epithelial Responses to SARS-CoV-2 Infection. *Nat Microbiol* **2024**, *9* (5), 1293–1311. https://doi.org/10.1038/s41564-024-01658-1.

(5) Loske, J.; Röhmel, J.; Lukassen, S.; Stricker, S.; Magalhães, V. G.; Liebig, J.; Chua, R. L.; Thürmann, L.; Messingschlager, M.; Seegebarth, A.; Timmermann, B.; Klages, S.; Ralser, M.; Sawitzki, B.; Sander, L. E.; Corman, V. M.; Conrad, C.; Laudi, S.; Binder, M.; Trump, S.; Eils, R.; Mall, M. A.; Lehmann, I. Pre-Activated Antiviral Innate Immunity in the Upper Airways Controls Early SARS-CoV-2 Infection in Children. *Nat Biotechnol* **2022**, *40* (3), 319–324. https://doi.org/10.1038/s41587-021-01037-9.

(6) Zhou-Suckow, Z.; Duerr, J.; Hagner, M.; Agrawal, R.; Mall, M. A. Airway Mucus, Inflammation and Remodeling: Emerging Links in the Pathogenesis of Chronic Lung Diseases. *Cell Tissue Res* **2017**, *367* (3), 537–550. https://doi.org/10.1007/s00441-016-2562-z.

(7) Whitsett, J. A. Airway Epithelial Differentiation and Mucociliary Clearance. *Ann Am Thorac Soc* **2018**, *15* (Suppl 3), S143–S148. https://doi.org/10.1513/AnnalsATS.201802-128AW.

(8) Kim, H.-T.; Yin, W.; Nakamichi, Y.; Panza, P.; Grohmann, B.; Buettner, C.; Guenther, S.; Ruppert, C.; Kobayashi, Y.; Guenther, A.; Stainier, D. Y. R. WNT/RYK Signaling Restricts Goblet Cell Differentiation during Lung Development and Repair. *Proceedings of the National Academy of Sciences* **2019**, *116* (51), 25697–25706. https://doi.org/10.1073/pnas.1911071116.

(9) Button, B.; Anderson, W. H.; Boucher, R. C. Mucus Hyperconcentration as a Unifying Aspect of the Chronic Bronchitic Phenotype. *Annals of the American Thoracic Society* **2016**.

(10) Rose, M. C.; Voynow, J. A. Respiratory Tract Mucin Genes and Mucin Glycoproteins in Health and Disease. *Physiol Rev* **2006**, *86* (1), 245–278. https://doi.org/10.1152/physrev.00010.2005.

(11) Chen, Y.; Hamati, E.; Lee, P.-K.; Lee, W.-M.; Wachi, S.; Schnurr, D.; Yagi, S.; Dolganov, G.; Boushey, H.; Avila, P.; Wu, R. Rhinovirus Induces Airway Epithelial Gene Expression through Double-Stranded RNA and IFN-Dependent Pathways. *Am J Respir Cell Mol Biol* **2006**, *34* (2), 192–203. https://doi.org/10.1165/rcmb.2004-0417OC.

(12) Schoggins, J. W.; Rice, C. M. Interferon-Stimulated Genes and Their Antiviral Effector Functions. *Current Opinion in Virology* **2011**, *1* (6), 519–525. https://doi.org/10.1016/j.coviro.2011.10.008.

(13) Nakajima, H.; Takatsu, K. Role of Cytokines in Allergic Airway Inflammation. *Int Arch Allergy Immunol* **2007**, *142* (4), 265–273. https://doi.org/10.1159/000097357.

(14) Saggini, A.; Maccauro, G.; Tripodi, D.; De Lutiis, M. A.; Conti, F.; Felaco, P.; Fulcheri, M.; Galzio, R.; Caraffa, A.; Antinolfi, P.; Felaco, M.; Pandolfi, F.; Sabatino, G.; Neri, G.; Shaik-Dasthagirisaheb, Y. B. Allergic Inflammation: Role of Cytokines with Special Emphasis on IL-4. *Int J Immunopathol Pharmacol* **2011**, *24* (2), 305–311. https://doi.org/10.1177/039463201102400204.

(15) Chung, F. Anti-Inflammatory Cytokines in Asthma and Allergy: Interleukin-10, Interleukin-12, Interferon-Gamma. *Mediators Inflamm* **2001**, *10* (2), 51–59. https://doi.org/10.1080/09629350120054518.

(16) Daines, S. M.; Orlandi, R. R. Inflammatory Cytokines in Allergy and Rhinosinusitis. *Curr Opin Otolaryngol Head Neck Surg* **2010**, *18* (3), 187–190. https://doi.org/10.1097/MOO.0b013e328338206a.

(17) L'Huillier, A. G.; Tapparel, C.; Turin, L.; Boquete-Suter, P.; Thomas, Y.; Kaiser, L. Survival of Rhinoviruses on Human Fingers. *Clin Microbiol Infect* **2015**, *21* (4), 381–385. https://doi.org/10.1016/j.cmi.2014.12.002.

(18) Geiser, J.; Boivin, G.; Huang, S.; Constant, S.; Kaiser, L.; Tapparel, C.; Essaidi-Laziosi, M. RSV and HMPV Infections in 3D Tissue Cultures: Mechanisms Involved in Virus-Host and Virus-Virus Interactions. *Viruses* **2021**, *13* (1), 139. https://doi.org/10.3390/v13010139.
